# Molecular basis of pathogenicity of the recently emerged FCoV-23 coronavirus

**DOI:** 10.1101/2024.08.25.607996

**Authors:** M. Alejandra Tortorici, Annette Choi, Cecily A. Gibson, Jimin Lee, Jack T. Brown, Cameron Stewart, Anshu Joshi, Sheri Harari, Isabelle Willoughby, Catherine Treichel, Elizabeth M. Leaf, Jesse D. Bloom, Neil P. King, Christine Tait-Burkard, Gary R. Whittaker, David Veesler

**Affiliations:** Department of Biochemistry, University of Washington, Seattle, Washington, USA; Department of Microbiology & Immunology, College of Veterinary Medicine, Cornell University, Ithaca, NY 14853, USA; Howard Hughes Medical Institute, Seattle, WA 98195, USA; Basic Sciences Division and Computational Biology Program, Fred Hutchinson Cancer Center; Institute for Protein Design, University of Washington, Seattle, WA 98195, USA Seattle, WA 98109, USA; The Roslin Institute, Royal (Dick) School of Veterinary Studies, University of Edinburgh, Easter Bush, Midlothian, United Kingdom; Public & Ecosystem Health, Cornell University, Ithaca, NY 14853, USA; Feline Health Center, Cornell University, Ithaca, NY 14853, USA

## Abstract

The ability of coronaviruses to recombine and cross species barriers affects human and animal health globally and is a pandemic threat. FCoV-23 is a recently emerged, highly pathogenic recombinant coronavirus responsible for a widespread outbreak of feline infectious peritonitis (FIP) likely linked to in-host viral evolution. Here, we report cryoEM structures of two FCoV-23 spike (S) isoforms explaining that the in-host loss of domain 0 observed in clinical samples enhances entry into cells and fusogenicity by facilitating protease access, leading to biotype switching and lethality. We show that FCoV-23 can use several aminopeptidase N (APN) orthologs as receptors and reveal the molecular determinants of receptor species tropism, including a glycan modulating human receptor utilization. We define antigenic relationships among alphacoronaviruses infecting humans and other mammalian species and identify a cross-reactive alphacoronavirus monoclonal antibody inhibiting FCoV-23 pseudovirus entry, paving the way for vaccine and therapeutic development targeting this highly pathogenic virus.

---

Coronaviruses circulating in wildlife pose a significant public health threat due to their zoonotic potential^1–4^. Additionally, cross-species transmission between animals can lead to the establishment of new disease reservoirs, impacting both wildlife populations and agricultural infrastructures^5,6^. Companion animals, such as cats and dogs, can play a role in these spillover events by serving as intermediate hosts not only for coronaviruses^7–9^ but also for influenza viruses^10,11^ and rabies virus^12^ for instance. The current expansion of highly pathogenic avian influenza H5N1 reservoirs in wild birds, poultry, marine mammals, and dairy cows, with occasional spillover to humans, highlights the ongoing risk of cross-species transmission^13,14^.

Alphacoronaviruses infect a wide range of animal species, and include the human pathogens HCoV-229E^15^, HCoV-NL63^16,17^ and the recently identified CCoV-HuPn-2018/HuCCoV_Z19Haiti^18–20^ as well as transmissible gastroenteritis virus (TGEV), porcine respiratory virus (PRCV), feline coronaviruses (FCoV) and canine coronaviruses (CCoV). Extensive recombination occurs among alphacoronaviruses taxonomically classified as a single virus species (*Alphacoronavirus-1*) but which infect several animal species, including cats, dogs, pigs and potentially rabbits^21–24^. Both serotypes of FCoV, designated FCoV-1 and FCoV-2, are major pathogens of wild felids and domestic cats capable of an in-host biotype switch from the relatively mild feline enteric coronavirus (FECV) to the highly pathogenic, macrophage-tropic feline infectious peritonitis virus (FIPV)^23,25–27^. Although the recent use of antiviral drugs in cats^28^ helped alleviate the mortality burden of FIP, this remains a prohibitively expensive option for many cat owners.

All coronaviruses are decorated with multiple copies of a homotrimeric S glycoprotein that is the target of neutralizing antibodies, drives viral entry through receptor binding and membrane fusion and contributes to pathogenesis^29–34^. Alphacoronaviruses were initially thought to all use aminopeptidase N (APN) as a primary receptor^35^ until the identification of ACE2 as the HCoV-NL63 receptor^36^ after the SARS-CoV-1 epidemic. Furthermore, the alphacoronavirus-specific S glycoprotein domain 0 mediates attachment to host cell surface sialosides, possibly facilitating the entry process^18,37–39^. Despite the diversity and medical/veterinary importance of these pathogens, cross-species transmission of animal alphacoronaviruses to humans and associated immunity remain largely under-studied.

In 2023, a novel FCoV-2 (named FCoV-23) was identified as the cause of a large outbreak of FIP in cats on the Mediterranean island of Cyprus, leading to documented spread of an import-related case to the UK^22,40,41^. The virus is highly virulent with most cats showing signs consistent with effusive FIP and profound neurological signs along with elevated viral loads in the colon in cells with macrophage-like morphology^22^. Genome sequencing suggested that FCoV-23 acquired its S gene and a small region of Orf1b from a pantropic, hypervirulent NA/09-like CCoV first identified in Greece and designated pCCoV^42,43^, with such viruses now circulating widely in Europe and possibly other parts of the world. Although the FCoV-23 genome is highly conserved among sequenced isolates, in-frame deletions of the S domain 0 occurred in an almost cat-specific manner in >90% of studied cases, suggesting in-host evolution^22^.

## Architecture of the FCoV-23 S glycoprotein

To unveil the 3D organization of the FCoV-23 infection machinery, we determined cryoEM structures of a prefusion-stabilized (using two prolines^44^) S long ectodomain trimer construct (**Fig. 1a,b, Extended Data Figure 1 and Table 1**). FCoV-23 S long comprises an N-terminal S_1_ subunit, folding as five domains designated, 0 and A to D, and a C-terminal S_2_ subunit (**Fig. 1a,b**). 3D classification of the cryoEM data revealed the presence of two distinct S conformations, with three D0 at the periphery of the trimer in a “swung out” conformation or with one D0 facing the viral membrane in a “proximal” conformation for which we determined structures at 2.3 Å and 2.7 Å resolution, respectively (**Fig. 1a,b and Extended Data Figure 1**). In both conformations, domain B (the receptor-binding domain) adopts a closed state (**Fig. 1a,b**). Moreover, we determined a 2.5 Å resolution cryoEM structure of a prefusion-stabilized FCoV-23 S ectodomain trimer lacking the N-terminal domain 0, representative of the in-host deletion occurring in >90% of sequenced viruses^22^. 3D classification of the cryoEM data revealed the presence of a single conformational state **(Extended Data Figure 2 and Table S1),** which is virtually identical to the long S structure besides the absence of domain 0. The overall FCoV-23 S architecture is reminiscent of that of CCoV-HuPn-2018 S with which it can be superimposed with an RMSD of 0.5 Å over 955 aligned Cα pairs. The S_1_ subunit folds with a ‘square-shaped’ tertiary structure, typical of α- and δ-coronavirus S glycoproteins, setting them apart from the ‘V-shaped’ organization observed for β- and ૪-coronaviruses^18,30,31,45–47^ (**Fig. 1c and Extended Data Figure 3**). The S_2_ subunit adopts a spring-loaded prefusion conformation and is closely similar to that of CCoV-HuPn-2018 with which it can be superimposed with an rmsd of 0.5 Å over 479 aligned Cα pairs (**Fig. 1d**). The absence of a polybasic cleavage site at the S_1_/S_2_ subunit junction concurs with the classification of FCoV-23 as part of FCoV-2^27^ and with the lack of proteolytic processing observed during S biogenesis, similar to CCoV-HuPn-2018 S^18^ (**Fig. 1g,h**).

**Table 1.** CryoEM data collection and refinement statistics.

| Data collection and processing | FCoV-23 S long<br>(swung-out D0)<br>Global refinement | FCoV-23 S long<br>(mixed D0)<br>Global refinement | FCoV-23 S<br>(swung-out D0)<br>Local refinement | FCoV-23 S<br>(proximal D0)<br>Local refinement | FCoV-23 S<br>short | fAPN + FCoV-23 RBD |
| --- | --- | --- | --- | --- | --- | --- |
| Magnification | 105,000 | 105,000 | 105,000 | 105,000 | 105,000 | 105,000 |
| Voltage (kV) | 300 | 300 | 300 | 105,000 | 105,000 | 105,000 |
| Electron exposure (e<sup>-</sup>/Å<sup>2</sup>) | 53.25 | 53.25 | 53.25 | 53.25 | 53.25 | 53.25 |
| Defocus range (-μm) | 0.8 - 1.7 | 0.8 - 1.7 | 0.8 - 1.7 | 0.8 - 1.7 | 0.8 - 1.7 | 0.8 - 1.7 |
| Pixel size (Å) | 0.829 | 0.829 | 0.829 | 0.829 | 0.829 | 0.829 |
| Symmetry imposed | C3 | C1 | C3 | C1 | C1 | C2 |
| Initial number of particles | 1,018,178 | 1,018,178 | 1,087,668 | 213,214 | 612,800 | 1,009,253 |
| Final number of particles | 362,667 | 213,214 | 539,331 | 117,776 | 262,922 | 466,950 |
| Map resolution (Å) | 2.3 | 2.7 | 2.4 | 3.2 | 2.5 | 2.5 |
| FSC threshold | 0.143 | 0.143 | 0.143 | 0.143 | 0.143 | 0.143 |
| Map sharpening B factor<br>(Å<sup>2</sup>) | -70.17 | -66.05 | -71.38 | -7.29 | -46.87 | -142.83 |
| **Validation** |  |  |  |  |  |  |
| MolProbity score | 1.07 | 0.79 | 1.59 | 1.17 | 1.1 | 1.26 |
| Clashscore | 1.53 | 0.62 | 6.96 | 1.46 | 1.83 | 3.95 |
| Poor rotamers (%) | 0.19 | 0.03 | 1.00 | 0.00 | 0.52 | 0.00 |
| **Ramachandran plot** |  |  |  |  |  |  |
| Favored (%) | 96.99 | 97.59 | 96.68 | 95.85 | 97.11 | 97.63 |
| Allowed (%) | 3.01 | 2.35 | 3.32 | 3.72 | 2.89 | 2.37 |
| Disallowed (%) | 0.00 | 0.06 | 0.00 | 0.41 | 0.00 | 0.00 |
| Entry codes | EMD<br>PDB | EMD<br>PDB | EMD<br>PDB | EMD<br>PDB | EMD<br>PDB | EMD<br>PDB |

Structure-based phylogenetic analysis clusters the FCoV-23 RBD with the CCoV-HuPn-2018, TGEV, PRCV, and FCoV-2 RBDs, underscoring their close evolutionary relationships (**Fig. 1e**). The similarity further extends to the conservation of key APN-interacting residues, including FCoV-23 S Y549/Q551 and W592 (CCoV-HuPN-2018 residues Y543/Q545 and W586)^18,48,49^, and to the conformational masking and glycan shielding of the receptor-binding loops by the N567 oligosaccharide (topological equivalent to CCoV-HuPn-2018 N561 glycan) from a neighboring RBD in the closed S trimer (**Fig. 1f**). FCoV-23 RBD mutations relative to other FCoV-2 sequences^22^ map at the periphery or outside the predicted APN-binding motif and are therefore not expected to affect host receptor tropism majorly. These findings suggest that FCoV-23 utilizes APN as an entry receptor through a similar binding mode to that observed for CCoV-HuPn-2018^18^ and PRCV/TGEV^50^. Strikingly, the FCoV-1 UU4 RBD is more distantly related to the FCoV-23 RBD than the α-coronavirus PEDV and HCoV-229E RBDs or porcine delta-coronavirus (PDCoV) RBD despite the classification of all FCoVs as a single viral species (**Fig. 1e**). This extensive genetic distance further concurs with the lack of APN utilization for FCoV-1, for which the receptor is unknown^48,51^, and for HCoV-NL63, which uses the ACE2 receptor ^36^.

**Figure 1.**
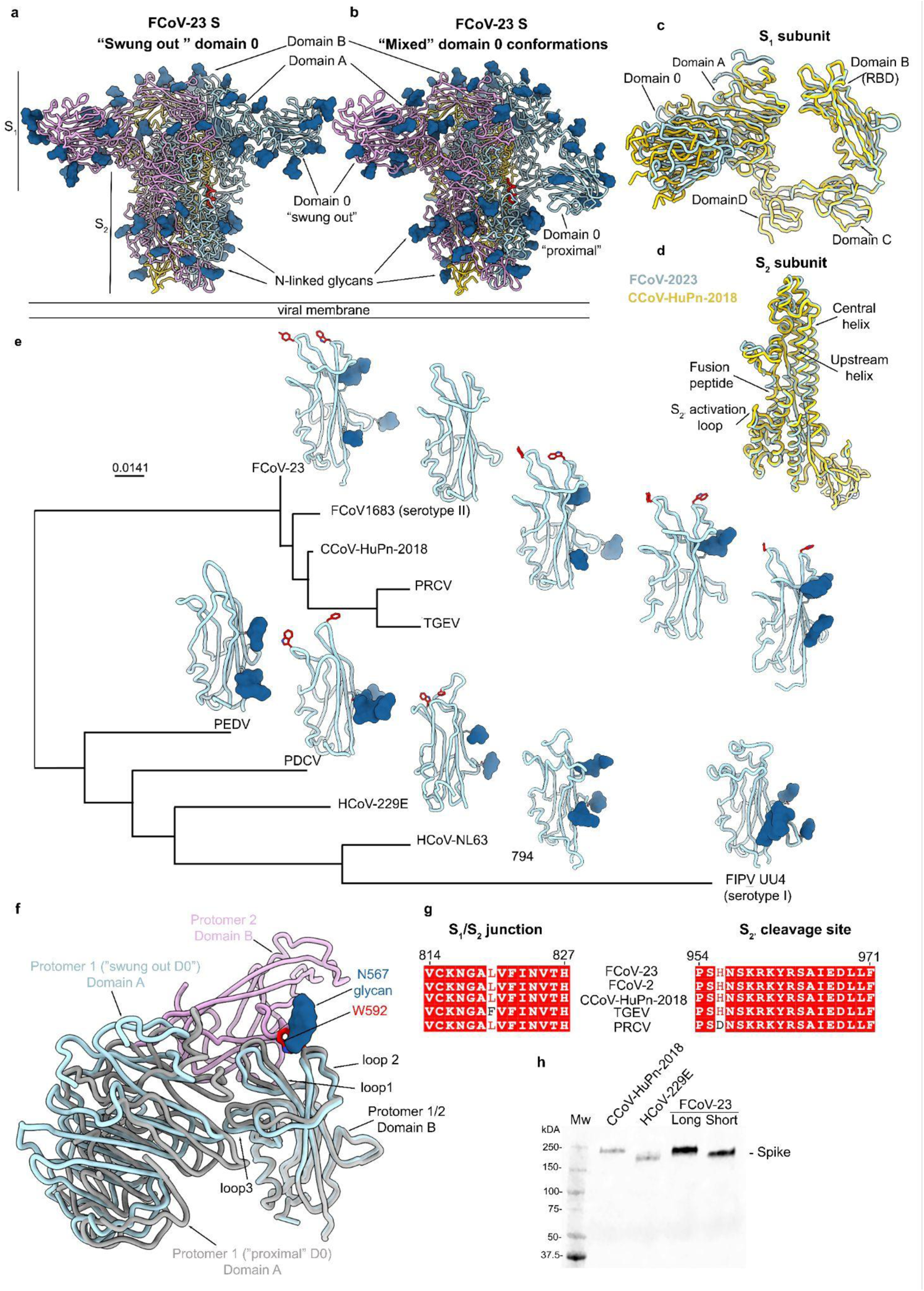
Architecture and evolution of the FCoV-23 S glycoprotein. **a,b**, Ribbon diagrams of the cryoEM structures of prefusion FCoV-23 S long in two distinct conformations defined by the positioning of domain 0. Each structure is rendered from a composite model of the global S refinement and local D0 refinement. The fusion peptide is rendered in red in the blue protomer to indicate its positioning relative to domain 0 in each conformation. **c,** Superimposition of FCoV-23 and CCoV-HuPn-2018 S1 subunits with swung out domain 0. **d**, Superposition of the FCoV-23 and CCoV-HuPn-2018 S2 subunits. **e,** Structure-based phylogenetic classification of α- and δ-coronavirus RBDs calculated with FoldTree^52^. Conserved APN-interacting residues are shown in red. **f,** Zoomed-in view of the domain A and RBDs (domain B) of the two FCoV-23 S conformations superimposed showing the conformational masking and glycan shielding of the receptor-binding loops in the context of the S trimer. **g,** Amino acid sequence conservation of the residues spanning the S1/S2 subunit junction and the S2’ site/fusion peptide region for CCoV-HuPn-2018 (A0A8E6CMP0), PRCV (Q84852), TGEV (Q0PKZ5) and FCoV-2 (J7F9W6). The residue numbering corresponds to FCoV-23 S. **h,** Western blot of VSV particles pseudotyped with FCoV-23 long and short S, HCoV-229E (AAK32191.1, P100E isolate) S and CCoV-HuPn-2018 S.

## Sensitivity of FCoV-23 S to host proteases

Host proteases are critical determinants of cell tropism, host range, and pathogenesis by activating coronavirus S glycoproteins for membrane fusion through cleavage^34,53,54^. To investigate the spectrum of host cell proteases that can possibly activate FCoV-23 S, we quantified the maximal velocity (Vmax) of cleavage of a fluorogenic peptide comprising residues _958_SKRKYR↓SAIE_967_ (↓ indicates the position of the predicted S_2’_ cleavage site scissile bond) in the presence of a panel of proteases previously involved in coronavirus S activation^55–59^. Trypsin, cathepsin L, plasmin, PC1 and factor Xa efficiently cleaved the FCoV-23 peptide whereas minimal processing was observed with furin (**Extended Data Fig. 4a**). Whereas trypsin is present in the digestive tract and could support enteric infections^45^, plasmin and factor Xa are proteases secreted systemically that are involved in the blood coagulation cascade and may promote the broad tissue tropism of FCoV-23 in cats. Indeed, plasmin cleavage and the presence of a tyrosine residue at position P2 of the influenza A/WSN/33 hemagglutinin cleavage site was previously associated with the neurotropism of this virus^60^, which could support the neurotropism of FCoV-23 in cats which also harbors a P2 tyrosine.

## CCoV-HuPn-2018 S, but not FCoV-23 S, hemagglutinates human erythrocytes

We previously showed that CCoV-HuPn-2018 S domain 0 hemagglutinates turkey, dog, and human erythrocytes in a sialic acid-dependent manner, suggesting a possible role of cell surface sialosides in viral entry^18^. To evaluate the ability of FCoV-23 S to engage cell-surface sialoside attachment factors, we incubated domain 0, A, or B multivalently displayed at the surface of the I53-50 nanoparticle (NP) with feline, chicken, bovine, turkey and human erythrocytes. CCoV-HuPn-2018 domain 0-NP (positive control) hemagglutinated human erythrocytes whereas we did not detect FCoV-23 S-mediated hemagglutination of any of the red blood cells tested **(Extended Data Figure 5a-f**). Although the FCoV-23 and CCoV-HuPn-2018 0 domains share only 26 % amino acid sequence identity, they can be superimposed with an rmsd of 2.3 Å over 190 aligned Cα pairs (out of 243 residues), underscoring their structural similarity. Nevertheless, structure-based sequence alignment of these two domains highlight numerous diverging regions that could explain the distinct hemagglutination phenotypes (**Extended Data Figure 5g-h**). These findings suggest that FCoV-23 S has a different specificity or lower binding affinity for cell surface sialosides, relative to CCoV-HuPn-2018, or that it does not engage sialoside attachment factors for entry.

## Aminopeptidase N is a receptor for FCoV-23

To test the possible APN utilization of the newly emerged FCoV-23, we assessed the ability of the FCoV-23 RBD to bind to several APN orthologs. We detected binding of dimeric feline and canine APNs, but not human or galline APNs, to the immobilized FCoV-23 RBD using biolayer interferometry (**Fig. 2a**). We determined binding avidities (_KD,app_) of 11 and 1.3 nM for feline and canine APNs, respectively (**Fig. 2b,c and Table 2**). We subsequently assessed the ability of HEK293T cells transiently transfected with a panel of APN orthologs to enable entry of VSV particles pseudotyped with FCoV-23 long and short S, CCoV-HuPn-2018 S or HCoV-229E S (**Fig. 2d,e and Extended Data Figure 6c,d**). Concurring with the binding data, feline and canine APNs promoted efficient entry of FCoV-23 S VSV into cells, as was the case for porcine APN (**Fig. 2d,e**). Galline APN enabled reduced but detectable entry whereas wild type human APN did not (**Fig. 2d,e**). These data show that several APN orthologs render cells susceptible to FCoV-23 S-mediated entry, irrespective of the presence or absence of domain 0. Finally, we observed concentration-dependent inhibition of FCoV-23 S long and short and CCoV-HuPn-2018 S VSV pseudovirus entry into cells mediated by feline and canine APN, underscoring the APN-specific entry pathway (**Fig. 2f,g and Extended Data Figure Fig 5e**). Collectively, these findings identify APN as a bona fide entry receptor for FCoV-23, which appears to have a broad receptor species tropism, mirroring that of CCoV-HuPn-2018.

**Table 2.** Biolayer interferometry kinetic parameters for binding of feline and canine APN to immobilized FCoV-23 RBD.

|  | Feline APN |  | Canine APN |  |
| --- | --- | --- | --- | --- |
|  | Replicate 1 | Replicate 2 | Replicate 1 | Replicate 2 |
| **K<sub>D,app</sub>** | 11 nM | 10 nM | 1.3 nM | 1.2 nM |
| **k<sub>on</sub>** | 1.2 · 10<sup>5</sup> M<sup>-1</sup>s<sup>-1</sup> | 1.3 · 10<sup>5</sup> M<sup>-1</sup>s<sup>-1</sup> | 1.5 · 10<sup>5</sup> M<sup>-1</sup>S<sup>-1</sup> | 1.5 · 10<sup>5</sup> M<sup>-1</sup>S<sup>-1</sup> |
| **k<sub>off</sub>** | 1.4 · 10<sup>-3</sup> s<sup>-1</sup> | 1.3 · 10<sup>-3</sup> s<sup>-1</sup> | 1.9 · 10<sup>-4</sup> S<sup>-1</sup> | 1.8 · 10<sup>-4</sup> S<sup>-1</sup> |

## Molecular basis of FCoV-23 utilization of aminopeptidase N

To understand APN recognition, we determined a cryoEM structure of the complex between the FCoV-23 RBD and feline APN at 2.5 Å resolution (**Fig. 2h,i, Extended Data Figure 6 and Table 1**). One FCoV-23 RBD binds to each of the two protomers of the feline APN dimer through interactions largely similar to those observed for the canine APN-bound CCoV-HuPn-2018 RBD^18^ and the porcine APN-bound PRCV RBD^50^ structures. The receptor-binding loop 1 Y549_FCoV-23_ side chain interacts with the N740_APN_ glycan core fucose via CH-π interactions^61^ and is hydrogen-bonded to the E735 and W741 side chains whereas the Q551_FCoV-23_ side chain is hydrogen bonded to the N740_APN_ side chain and interacting with the proximal N-acetyl-glucosamine and core fucose (**Fig. 2i**). In receptor-binding loop 2, W592 is packed against APN residues H790_APN_ and P791_APN_ and its imino group is hydrogen-bonded to the main-chain carbonyl of N787_APN_. We note that the substitution of R540_CcoV-HuPn-2018_ with L546_FCoV-23_ abrogates a salt bridge formed with the canine E786_APN_ corresponding to feline Q779_APN_.

Given the similarities of the FCoV-23 and CcoV-HuPn-2018 RBDs and of their host receptor tropism, we hypothesized that the lack of an oligosaccharide at position N739 of human APN, which is present in feline, canine, porcine and galline APNs, explained the inability of FCoV-23 to engage the human APN ortholog, as is the case for CCoV-HuPn-2018^18^. Indeed, the human APN N739 oligosaccharide knockin mutant (R741T substitution), rendered cells permissive to FCoV-23 S VSV, confirming the importance of this glycan-mediated interaction for receptor engagement (**Fig. 2d,e**). Conversely, removal of this oligosaccharide from feline APN (T742R substitution) reduced FCoV-23 S-and CCoV-HuPn-2018 S-mediated entry into cells by two orders of magnitude (**Fig. 2d,e and Extended Data Figure 7c**), further demonstrating its key role for binding. Efficient HCoV-229E S-mediated entry occurred with human, feline and galline APNs and did not depend on the presence of a glycan at positions N739 (human APN) or N740 (feline APN) **(Extended Data Figure 7d**), concurring with the fact that the HCoV-229E RBD recognizes a different APN site relative to the other viruses evaluated here^62^.

**Figure 2.**
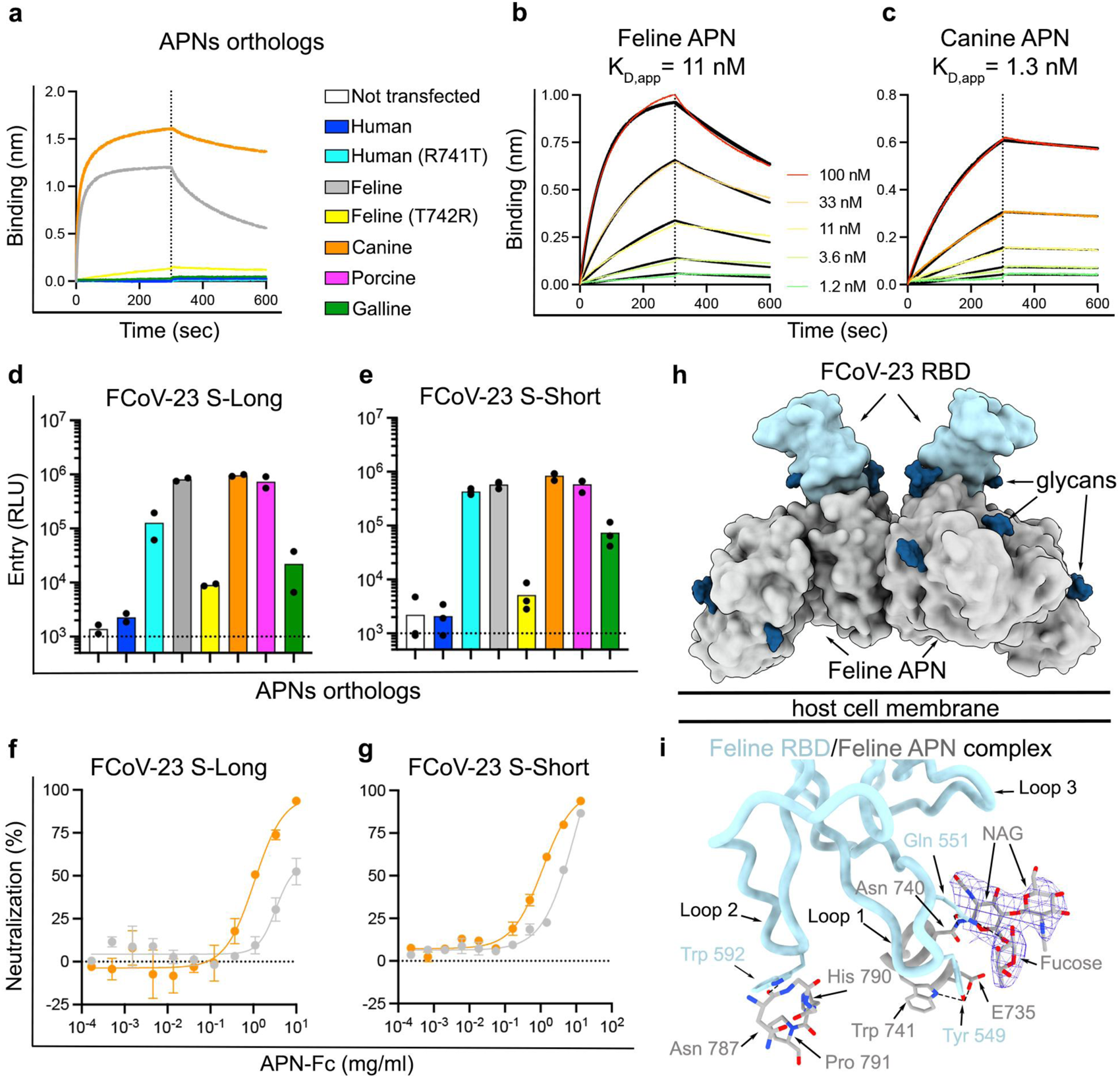
APN is a functional entry receptor for FCoV-23. **a**, Biolayer interferometry binding analysis of dimeric human, human R741T (glycan knockin), feline, feline T742R (glycan knockout), canine and galline APN-Fc ectodomains at a concentration of 1 µM to biotinylated FCoV-23 RBD immobilized at the surface of SA biosensors. **b,c,** Biolayer interferometry binding kinetic analysis of the indicated concentrations of dimeric feline and canine APN-Fc ectodomains to biotinylated FCoV-23 RBD immobilized on SA biosensors. Global fit (1:1 model) to the data is shown in black and reported affinities are expressed as apparent KD (KD, app) due to avidity resulting from the dimeric nature of APN. A representative experiment is shown out of two biological replicates. **d,e,** Entry of VSV particles pseudotyped with FCoV-23 S long (**d**) and FCoV-23 S short (**e**) into HEK293T cells transiently transfected with membrane-anchored feline, feline T742R (glycan knockout), canine, porcine, galline, human and human R741T (glycan knockin) APN orthologs. RLUs, relative luciferase units. Each dot represents a biological experiment each performed with technical duplicates or triplicates. **f,g**, Concentration-dependent inhibition mediated by purified dimeric feline and canine APN-Fc ectodomains of FCoV-2023 S long (**f**) and FCoV-23 short (**g**) S VSV pseudotyped virus entry into HEK293T cells transiently transfected with membrane-anchored feline APN. Each curve represents a biological experiment performed with technical duplicates. Error bars represent the standard error of the mean (SEM). **h,** Surface representation of the cryoEM structure of the FCoV-23 RBD (light blue) in complex with the feline APN ectodomain dimer (gray) depicted in its likely orientation relative to the host plasma membrane. N-linked glycans are rendered as dark blue surfaces. **i,** Close-up view showing key interactions formed between the FCoV-23 RBD (light blue) and feline APN (gray). Dashed lines show salt bridges and hydrogen bonds. The cryoEM density around the N740 glycan is shown as a blue mesh.

## FCoV-23 S is tropic for feline and canine cells

To gain insight into the cell tropism of FCoV-23, we pseudotyped MLV particles with FCoV-23 short and long S constructs and assessed their infectivity using a panel of cell lines representing multiple mammalian species and tissues. Both FCoV-23 short and long S MLV pseudoviruses effectively infected Fcwf-CU feline macrophage cells, CRFK feline kidney epithelial cells and AK-D feline epithelial lung cells with a consistently higher entry of short S MLV, relative to long S MLV (mean: 1.5-3, **Fig. 3a,b and Extended Data Figure 4c**). Furthermore, we detected weak pseudovirus entry into canine A-72 fibroblast cells with FCoV-23 short S MLV, but not FCoV-23 long S MLV, further supporting the superior functionality of the former construct (**Fig. 3a,b and Extended Data Figure 4c**).

## Domain 0 deletion enhances FCoV-23 S fusogenicity

To extend these findings and assess the ability of FCoV-23 S to induce syncytia formation, we performed a cell-cell fusion assay by transiently transfecting either the FCoV-23 short or the long S construct in the aforementioned panel of cell lines. In line with the pseudovirus entry data, syncytia were observed with Fcwf-CU, CRFK, AK-D feline cells and with A72 canine cells whereas minimal to no fusion could be detected in the other cells tested (**Fig. 3c and Extended Data Figure 8**). FCoV-23 short S induced larger syncytia and with faster kinetics than FCoV-23 long S pointing to higher fusogenicity of the domain 0-lacking S construct, mirroring the higher MLV pseudovirus infectivity in feline cells. For instance, large syncytia appeared as early as four hours post-transfection for FCoV-23 short S, but not for long S, and transfection in Fcwf-CU cells led to cell lifting by twelve hours (**Fig. 3c**). Collectively, these data suggest that FCoV-23 has a broad feline cell and tissue tropism, with limited to no permissiveness for other cells tested here, concurring with the pathology observed in FCoV-23-infected cats^22^.

**Figure 3.**
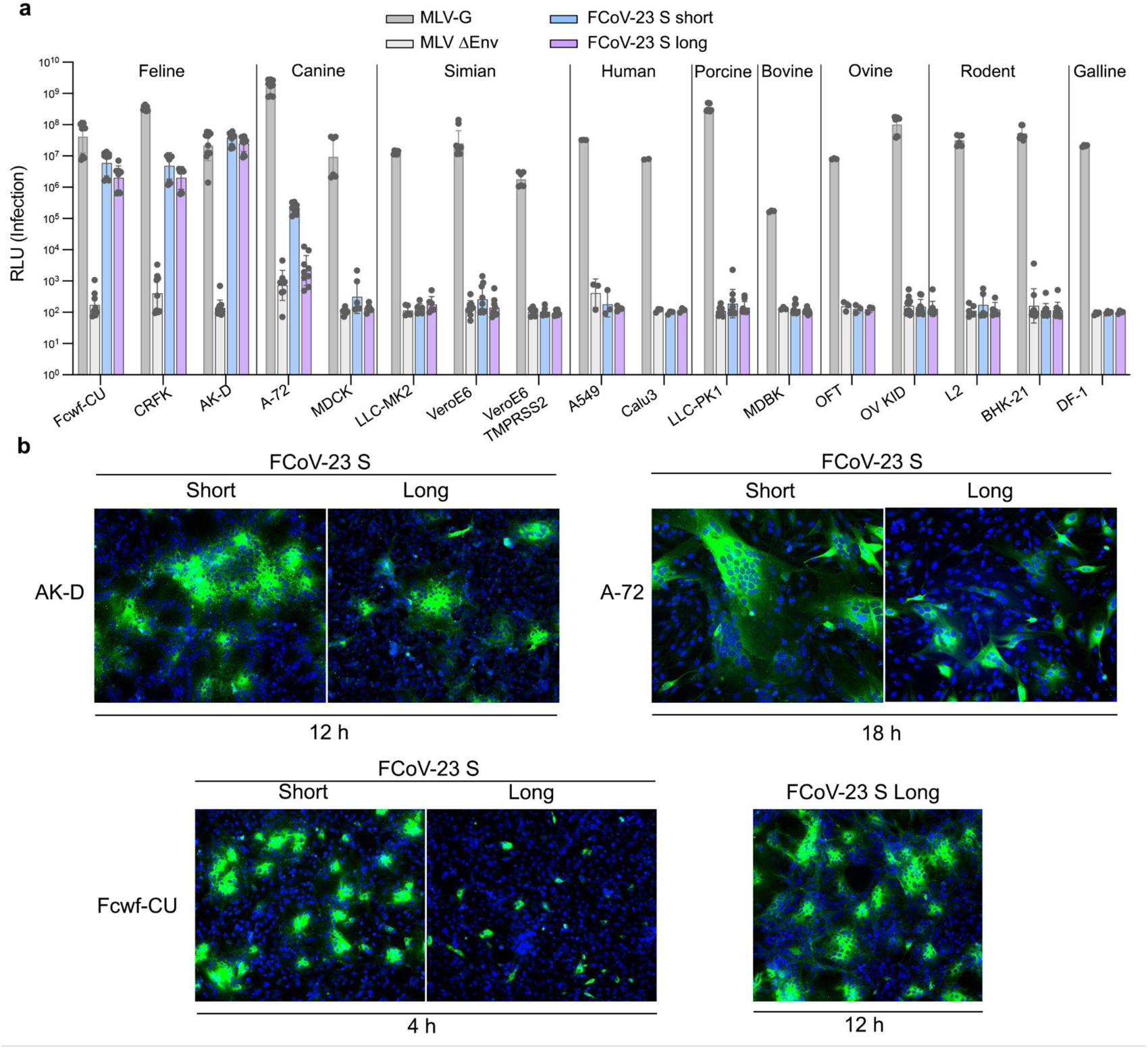
FCoV-23 has a broad feline cell and tissue tropism. **a**, Entry of MLV particles pseudotyped with FCoV-23 S long and short in a panel of cell lines spanning various species and tissues as follows. Feline: Fcwf-CU (feline macrophage-like cells), CRFK (feline epithelial kidney cells), AK-D (feline airway epithelial cells). Canine: A-72 (canine tumor fibroblast cells), MDCK (canine epithelial kidney cells). Simian: LLC-MK2 (Rhesus monkey kidney cells), Vero-E6 (African green monkey kidney cells), Vero-E6-TMPRSS2 (VeroE6 cells with stable TMPRSS2 expression). Human: A549 (human lung carcinoma epithelial cells) and Calu3 (human lung adenocarcinoma epithelial cells). Porcine: LLC-PK1 (porcine epithelial kidney cells). Bovine: MDBK (bovine epithelial kidney cells). Ovine: OFT (ovine fetal turbinate cells), OV KID (ovine kidney cells). Rodent: L2 (rat lung epithelial cells), BHK-21 (Syrian golden hamster kidney cells). MLV-G: MLV particles expressing VSV-G glycoprotein were used as a positive control. MLV ΔEnv: MLV particles with no envelope were used as negative control. RLU: Relative luciferase unit. Bars represent the geometric mean obtained from nine technical replicates. **b,** Cell-Cell fusion assay in AK-D, A-72 and Fcwf-Cu cells expressing FCoV-23 S long and short. Cells were fixed at the indicated times and FCoV-23 S was detected using HA antibody (green) and nuclei were detected using DAPI stain (blue).

## Antigenicity of the FCoV-23 S glycoprotein

FCoV-23 S shares ∼82.11% and ∼48.2% amino acid sequence identity with CCoV-HuPN-2018 S and HCoV-229E S (AAK32191.1, P100E isolate), respectively, with the S_1_ subunits being more divergent than the S_2_ subunits. Furthermore, the FCoV-23 RBD shares ∼93.5% and ∼27.5% amino acid sequence identity with the CCoV-HuPN-2018 and HCoV-229E RBDs, respectively.

To investigate the antigenic relationships among alphacoronaviruses, we first evaluated the ability of the PRCV/TGEV RBD-directed 1AF10 neutralizing monoclonal antibody to inhibit FCoV-23 S-mediated entry. We found here that 1AF10 Fab neutralized FCoV-23 S VSV pseudovirus with similar potency to that observed against CCoV-HuPn-2018 S^18^ (**Fig. 4a,b and Extended Data Figure 6f-g**). These findings support the antigenic relationships between FCoV-23 S, CCoV-HuPn-2018, PRCV and TGEV S, which all cross-react with this monoclonal antibody blocking receptor engagement, identifying 1AF10 as a possible countermeasure against these viruses.

We investigated the neutralizing activity of human plasma samples collected in 2020 that neutralize HCoV-229E infection (**Table 3**) of mouse hyperimmune serum obtained upon vaccination with prefusion HCoV-229E S 2P (AAK32191.1, P100E isolate, 2001). Some of these plasma samples weakly reduced FCoV-23 S VSV and CCoV-HuPn-2018 S VSV entry into cells and their cross-neutralization potency was not necessarily correlated with that observed against HCoV-229E (**Fig. 4c,f**). Furthermore, although all mouse sera completely neutralized HCoV-229E S VSV pseudovirus, they exhibited weak to no inhibition of FCoV-23 S VSV and CCoV-HuPn-2018 S VSV (**Fig. 4g,i**). These data show that prior HCoV-229E infection or vaccination would elicit (at best) weak cross-neutralization of FCoV-23 S VSV and CCoV-HuPn-2018 S VSV, in line with the more distant genetic relationships between the S glycoprotein, and in particular RBDs, of HCoV-229E and these viruses.

**Figure 4.**
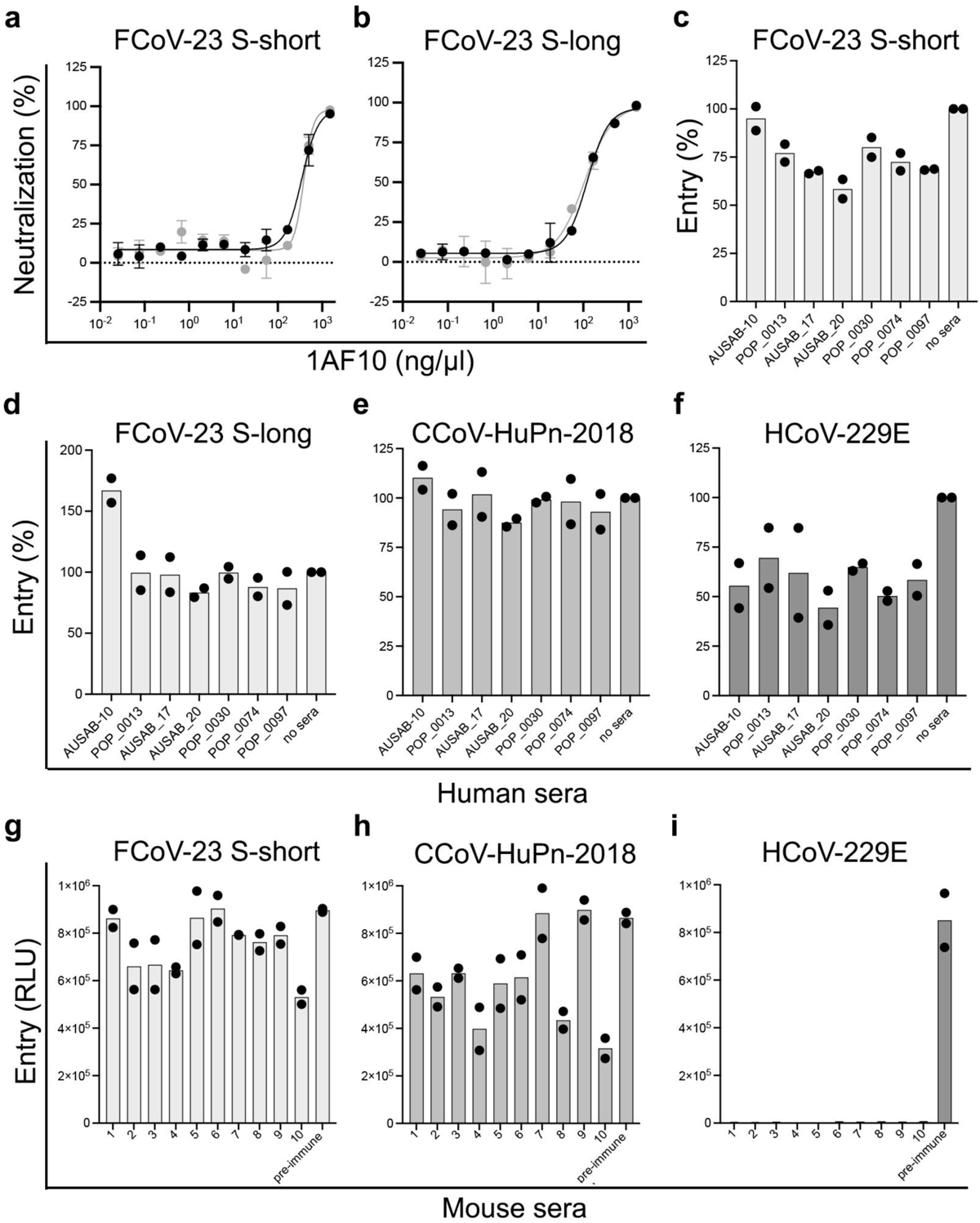
Cross-neutralization of FCoV-23 by α-coronavirus-elicited antibodies. **a,b** Dose-dependent neutralization of FCoV-23 S long (**a**) and FCoV-23 S short (**b**) pseudotyped VSV in the presence of various concentrations of the 1AF10 neutralizing monoclonal Fab fragment using HEK293T cells transiently transfected with membrane-anchored feline APN. **c-e,** Inhibition of FCoV-23 S long (**c**), FCoV-23 S short (**d**) CCoV-HuPn-2018 S (**e**) and HCoV-229E S, AAK32191.1, P100E isolate 2001 (**f**) VSV pseudovirus entry into HEK293T cells transiently transfected with membrane-anchored feline APN (FCoV-23, **c,d**), canine APN (CCoV-HuPn-2018, **e**) and human APN (HCoV-229E, **f**) in the presence of a 1/10 dilution of a panel of human sera with neutralizing activity against HCoV-229E. Two biological replicates each performed with technical duplicates is shown. **g,i,** Inhibition of FCoV-23 S short (**g**) CCoV-HuPn-2018 S (**h**) and HCoV-229E S, AAK32191.1, P100E isolate, 2001 (**i**) VSV pseudovirus entry into HEK293T cells transiently transfected with membrane-anchored feline APN (FCoV-23, **g**), canine APN (CCoV-HuPn-2018, **h**) and human APN (HCoV-229E, **i**) in the presence of a 1/10 dilution of HCoV-229E-S (AAK32191.1, P100E isolate, 2001) elicited mice sera.

**Table 3.**
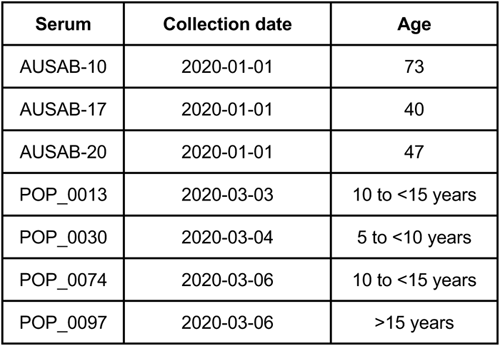
Demographics information of human serum samples.

| Serum | Collection date | Age |
| --- | --- | --- |
| AUSAB-10 | 2020-01-01 | 73 |
| AUSAB-17 | 2020-01-01 | 40 |
| AUSAB-20 | 2020-01-01 | 47 |
| POP_0013 | 2020-03-03 | 10 to <15 years |
| POP_0030 | 2020-03-04 | 5 to <10 years |
| POP_0074 | 2020-03-06 | 10 to <15 years |
| POP_0097 | 2020-03-06 | >15 years |

## Discussion

Most alphacoronavirus S glycoproteins harbor a domain 0, not present in other coronavirus genera, that arose through duplication of domain A (the latter domain is often referred to as the N-terminal domain for betacoronaviruses)^47^. Both domains adopt a similar galectin-like β-sandwich fold and have been shown to interact with host cell sialoside attachment factors for alphacoronavirus CCoV-HuPn-2018^18^, TGEV^64^, PEDV^38,39^ 0 domains and for OC43/HKU1^65–67^, MERS-CoV^68^ and SARS-CoV-2^69^ A domains. The FCoV-23 S structures presented here reveal distinct domain 0 conformations, akin to that observed for CCoV-HuPn-2018^18^ and PEDV^70–72^, likely corresponding to snapshots of the conformational changes leading to viral entry into cells. Strikingly, in-frame deletions of the FCoV-23 S domain 0 occurred in an almost cat-specific manner in >90% of studied Cypriot and import FIP cases, suggesting in-host evolution^22^, and is reminiscent of the evolutionary process that led from TGEV to PRCV^64,73^. The high pathogenicity of FCoV-23 and its detection in macrophage-like cells by immunohistochemistry are characteristics of the FECV to FIPV biotype switch. Furthermore, the markedly greater fusogenicity of FCoV-23 S short observed in various cell lines, relative to the long construct, suggests that deletion of domain 0 may promote increased syncytia formation and pathology, as described for the SARS-CoV-2 Delta variant as a result of the P681R substitution^74^. Our structural data reveal that the proximal domain 0 conformation would sterically limit access of activating host proteases to the fusion peptide and S_2’_ cleavage site. This offers a possible explanation for the repeated loss of domain 0 among FCoV-23-infected cats and delineating how domain 0 conformational changes may couple host cell attachment with membrane fusion to initiate alphacoronavirus infection.

Coronavirus RBDs are main targets of neutralizing antibodies and typically account for most plasma neutralizing activity elicited by infection or vaccination against matched and mismatched viruses^62,63,75–79^. Structure-guided phylogenetic classification clusters the FCoV-23, CCoV-HuPn-2018^18^, PRCV/TGEV^50^ and FCoV-2 RBDs, which engage the APN entry receptor with a similar binding mode. This clustering is further supported by the observation that FCoV-23 S promotes membrane fusion with and entry into Fcwf-CU feline macrophage-like cells, a property shared with FCoV-2 but not FCoV-1, the latter group of viruses using a distinct (yet unknown) receptor due to its markedly distinct receptor-binding loops^48,51,80^. The observation that the FCoV-1 RBD architecture^80^ is more divergent from the FCoV-2 RBD than from the HCoV-229E^62^ and PEDV^71^ RBDs, underscores the distinct receptor usage and evolutionary pathways of these two types of feline pathogens which will likely require the development of serotype-specific vaccines. The identification of APN as the FCoV-23 entry receptor along with that of a broad spectrum of proteases supporting proteolytic S processing illuminate key factors modulating host and cell tropism and cross-species transmission. The rare detection of weak cross-neutralization observed with a few 229E-elicited sera underscores the antigenic distance between this human endemic coronavirus and FCoV-23 although both pathogens belong to the alphacoronavirus genus. However, some degree of broad humoral immune protection may result from alphacoronavirus exposure, either through direct viral neutralization of distinct alphacoronaviruses or antibody-mediated Fc-effector functions, the latter mechanism was observed with S-elicited fusion machinery-directed polyclonal antibodies upon mismatched sarbecovirus challenge^81^.

Cross-species transmission of pathogens is one of the greatest threats to human and animal health worldwide and is exacerbated by travel, urbanization, deforestation and intensive farming. This is currently illustrated by the spillover of highly pathogenic avian influenza H5N1 virus to birds and mammals, including dairy cattle, cats and humans^13,14^. The *Alphacoronavirus-1* species comprise a set of highly recombinogenic viruses circulating in dogs, cats and pigs which has been proposed to be divided in two clades with functionally distinct S glycoproteins^21,23^. Furthermore, several *Alphacoronavirus-1* recombinant viruses have spilled over from dogs or cats to humans^9^. The wide circulation of *Alphacoronavirus-1* combined with our poor understanding of their genetic diversity and the accelerating rate of viral spillovers to the human population emphasizes the importance of preparing for the emergence of highly pathogenic coronaviruses.

## Acknowledgements

This study was supported by the National Institute of Allergy and Infectious Diseases (P01AI167966 to N.P.K, J.D.B. and D.V., DP1AI158186 and 75N93022C00036 to D.V.), an Investigators in the Pathogenesis of Infectious Disease Awards from the Burroughs Wellcome Fund (D.V.), the University of Washington Arnold and Mabel Beckman cryoEM center, the National Institute of Health grant S10OD032290 (to D.V.), the Morris Animal Foundation (G.W.), EveryCat Health Foundation (G.W.) and the Cornell Feline Health Center (G.W.). J.D.B. and D.V. are Investigators of the Howard Hughes Medical Institute and D.V. is the Hans Neurath Endowed Chair in Biochemistry at the University of Washington. C.T.B. is supported by the BBSRC Institute Strategic Programme grant funding to the Roslin Institute, grant numbers BBS/E/D/20241866, BBS/E/D/20002172, and BBS/E/D/20002174. AC is supported by a Liz Hanson Scholarship in Feline Medicine at Cornell University College of Veterinary Medicine.

## Author Contributions

M.A.T. and D.V. designed the study and the experiments; M.A.T. designed the constructs used in this study and M.A.T., C.G., J.B. and C.S. recombinantly expressed and purified glycoproteins. C.G. performed binding assays. M.A.T. and A.C. carried out pseudovirus entry assays. M.A.T. performed pseudovirus neutralization assays. M.A.T. and C.G. assembled nanoparticles and M.A.T. conducted hemagglutination assays. M.A.T. carried out cryoEM specimen preparation, data collection and processing. M.A.T. and D.V. built and refined atomic models. A.C. conducted fusion and cleavage assays under the supervision of G.W. M.A.T. and A.C. ran Western blots. E.M.L., I.W., and C.T. conducted the immunogenicity study and C.T.B. contributed unique reagents or data. M.A.T., A.C., C.G., J.L., G.W. and D.V. analyzed the data. M.A.T and D.V. wrote the manuscript with input from all authors; D.V. supervised the project.

---

**Extended Data Figure 1.**
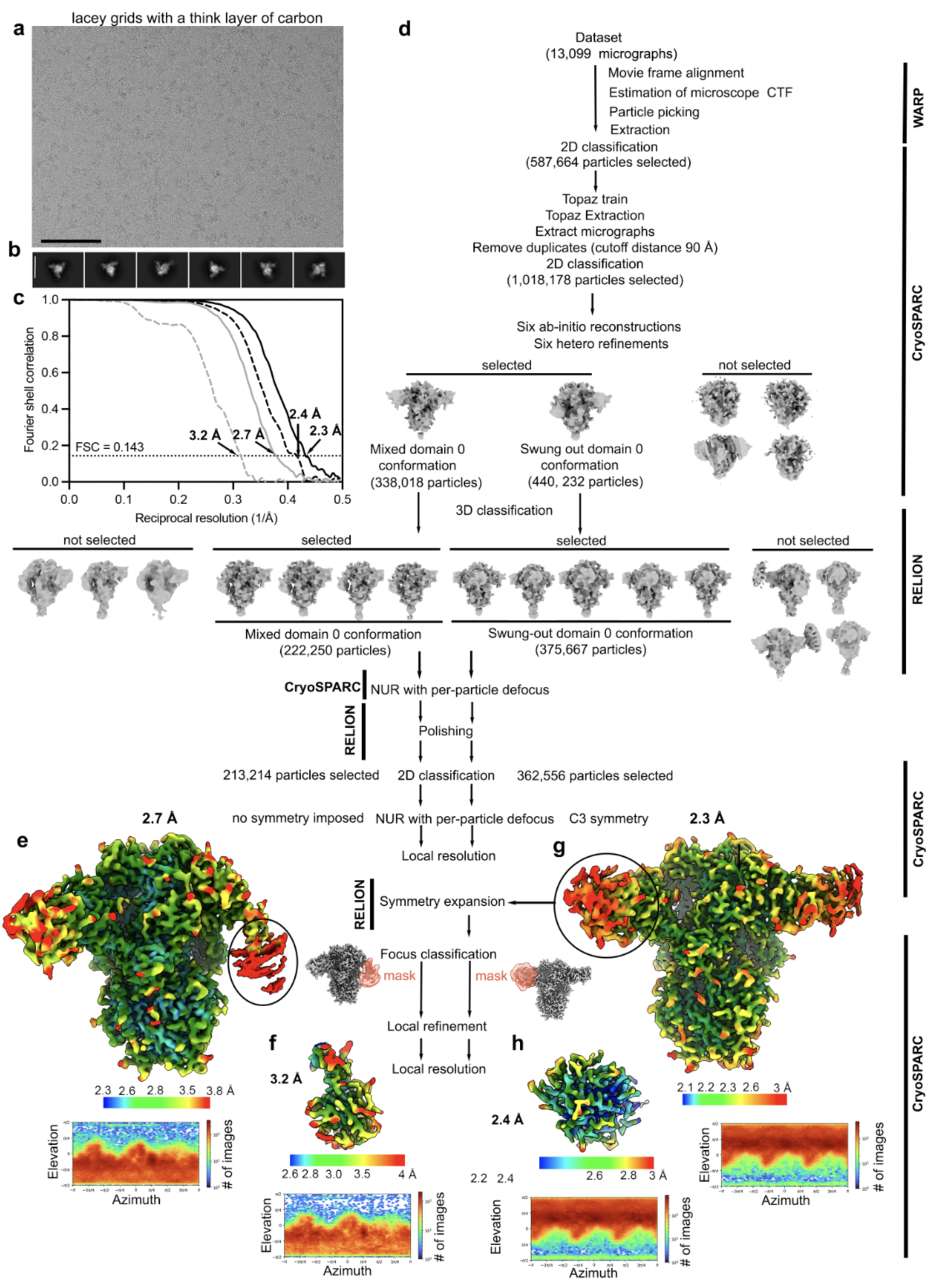
Data processing and validation of the FCoV-23 S long glycoprotein (with domain 0) dataset, related to Figures 1 and 2. **a,b,** Representative electron micrograph (**a**) and 2D class averages (**b**) of the prefusion FCoV-23 S long ectodomain trimer embedded in vitreous ice. Scale bar of the micrograph and the 2D class averages, 100 Å. **c,** Gold-standard Fourier shell correlation (FSC) curves for the FCoV-23 S trimer with domain 0 in the “swung out” conformation (black line), the FCoV-23 S trimer with mixed domain 0 conformation (gray line), the local refinements of domain 0 in swung out conformation (dashed black line) and in the proximal conformation (dashed gray line). The 0.143 cutoff is indicated by a horizontal dotted black line. **d,** Cryo-EM data processing flowchart. **e,f,** Unsharpened map of the FCoV-23 S trimer with mixed domain 0 conformations (**e**) and the locally refined unsharpened map of the proximal domain 0 (**f**) colored according to local resolution determined using cryoSPARC resolution. **g,h,** Unsharpened map of the FCoV-23 S trimer with domain 0 in swung out conformation (**g**) and the locally refined unsharpened map of a swung out domain 0 (**h**) colored according to local resolution determined using cryoSPARC resolution. Angular distribution plots with all the particles contributing to the final maps are shown at the bottom of each corresponding map. CTF: contrast transfer function. NUR: non uniform refinement.

**Extended Data Figure 2.**
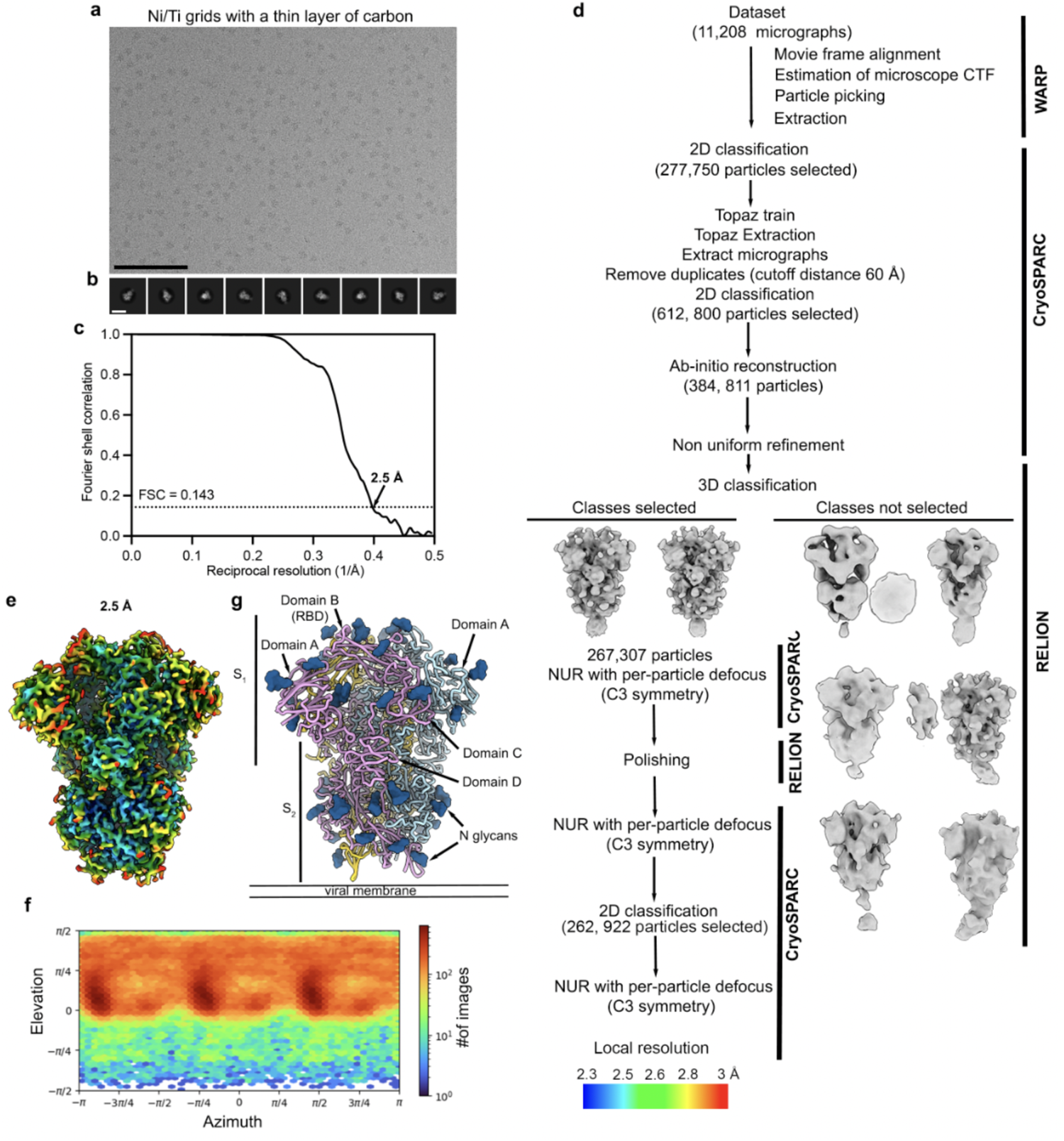
Data processing and validation of the FCoV-23 S glycoprotein (without domain 0) dataset, related to Figures 1 and 2. **a,b,** Representative electron micrograph (**a**) and 2D class averages (**b**) of the prefusion FCoV-23 S short (without domain 0) ectodomain trimer embedded in vitreous ice. Scale bar of the micrograph and the 2D class averages, 100 Å. **c,** Gold-standard Fourier shell correlation curve for the FCoV-23 S trimer reconstruction. The 0.143 cutoff is indicated by a horizontal dotted black line. **d,** Cryo-EM data processing flowchart. **e,** Unsharpened map of the FCoV-23 S short ectodomain trimer colored according to local resolution determined using cryoSPARC. **f,** Angular distribution plot of all the particles that contributed to the map. **g,** Ribbon representation of FCoV-23 S short highlighting the S1 and S2 subunits. N-linked glycans are shown in surface representation rendered in dark blue. CTF: contrast transfer function. NUR: non uniform refinement.

**Extended Data Figure 3.**
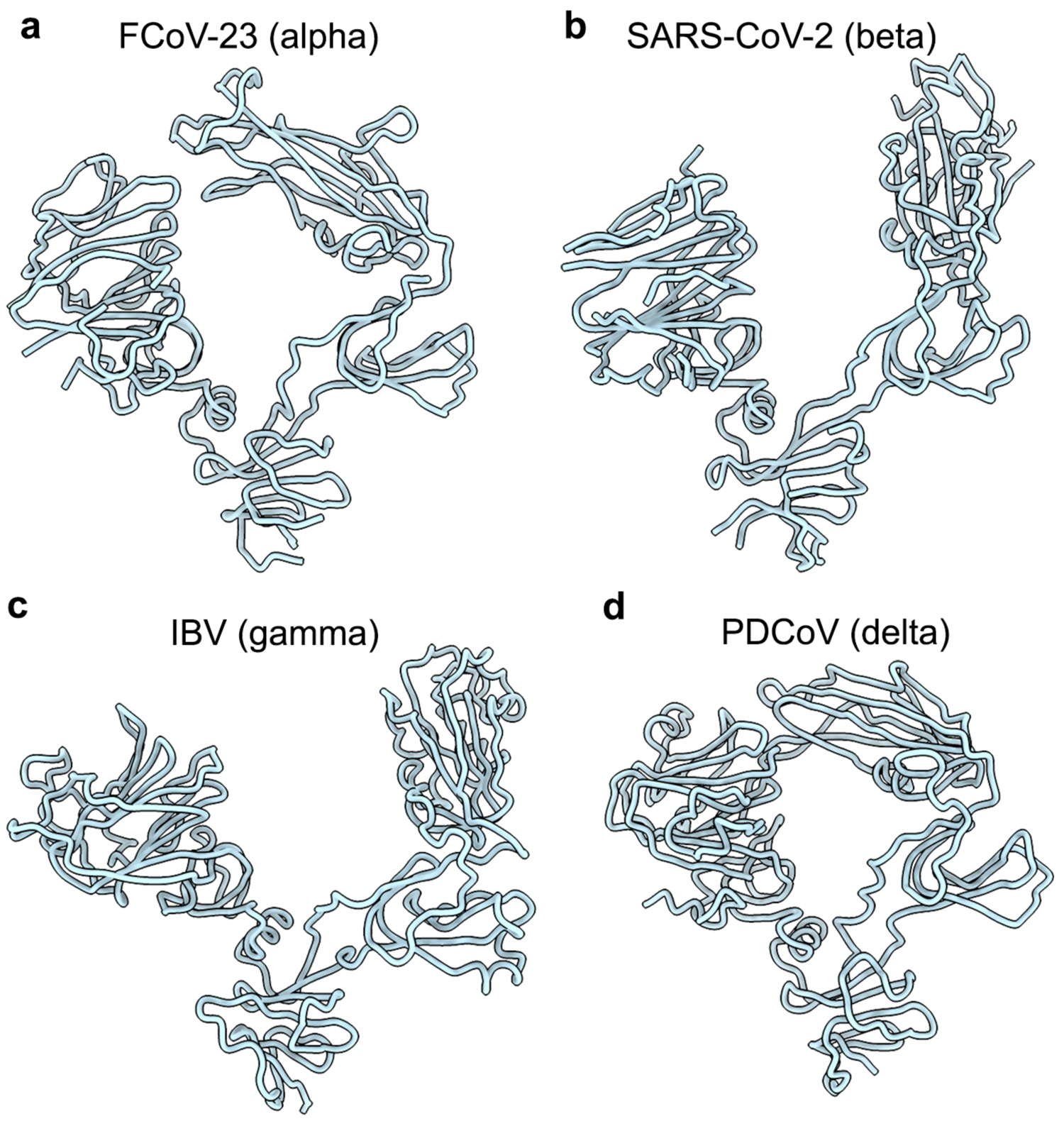
S1 subunit architecture of the S glycoprotein of all four coronavirus genera. **a-d**, Ribbon diagram of the FCoV-23 S short (**a**), SARS-CoV-2 (PDB 6VXX^45^, **b**), IBV (PDB 6CV0^46^ , **c**) and PDCoV (PDB 6BFU^45^, **d**) S1 subunits.

**Extended Data Figure 4.**
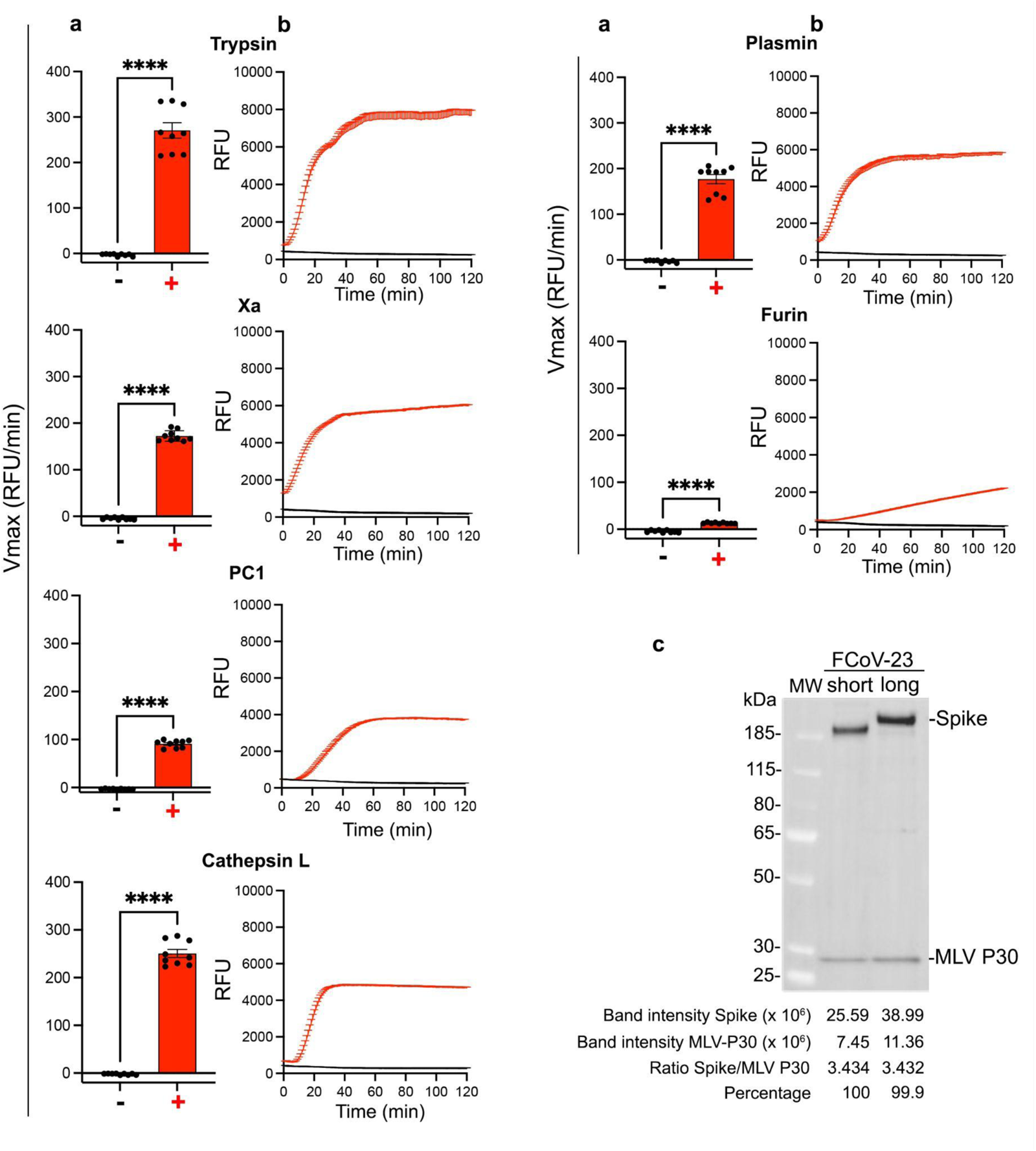
Sensitivity of FCoV-23 S to host proteases. **a,** Quantification of the maximal velocity (Vmax) of cleavage of the fluorogenic FCoV-23 S2’ peptide by the indicated proteases. The S2’ peptide was coupled to a fluorescence resonance energy transfer pair of fluorophores: 7-methoxycoumarin-4-yl acetyl (MCA) at the N-terminus and N-2,4-dinitrophenyl (DNP) at the C-terminus yielding MCA-958SKRKYRSAIE967-DNP. Dots represent technical replicates and bars correspond to the mean. Statistical analysis was performed using unpaired T test **** P < 0.0001. **b,** Cleavage of the fluorogenic FCoV-23 S2’ peptide, expressed in relative fluorescence intensity (RFU) as a function of time, by the indicated proteases. **c,** Quantification of FCoV-23 short and long S in MLV pseudotyped particles.

**Extended Data Figure 5.**
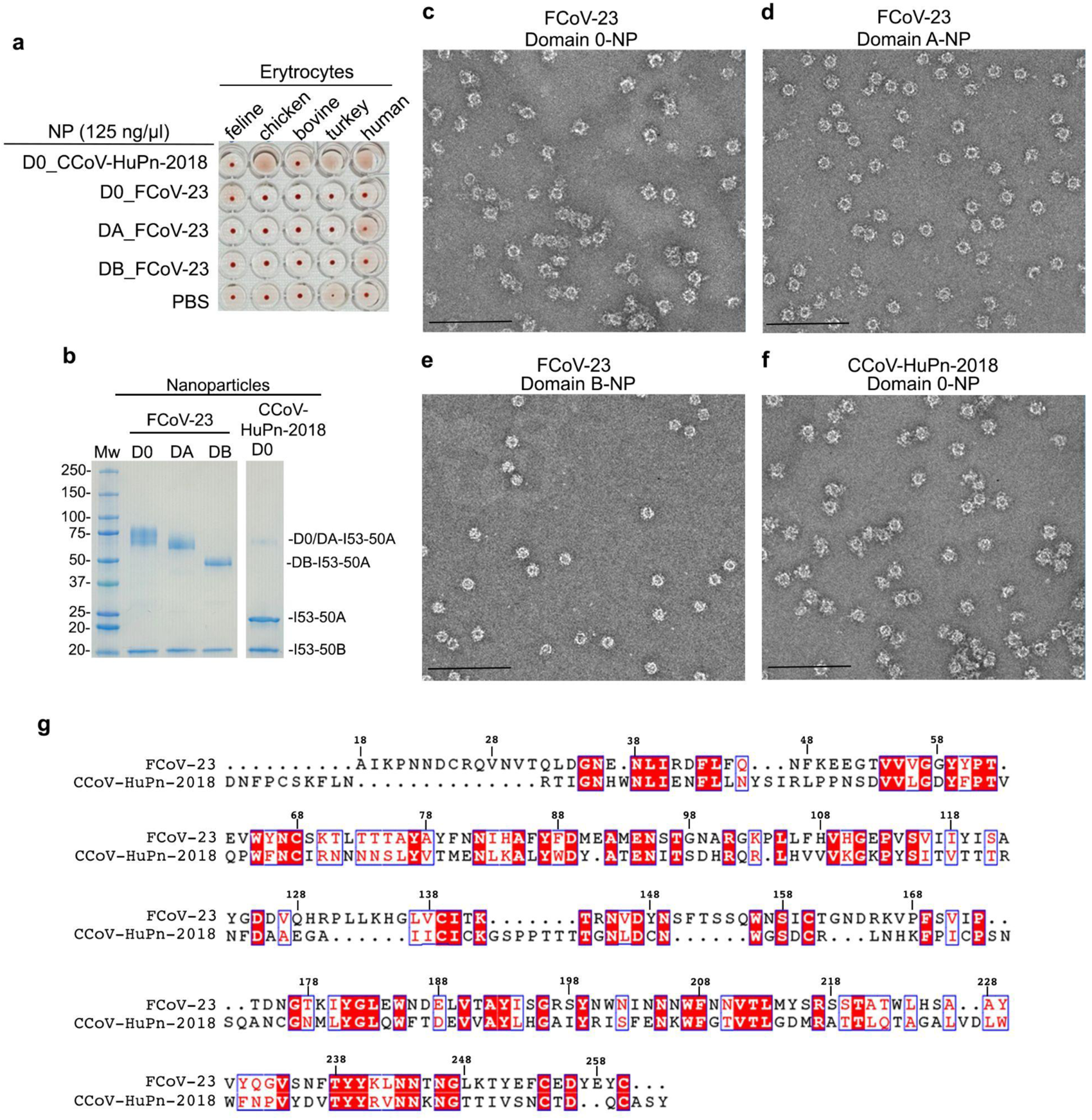
FCoV-23 S domain 0 does not hemagglutinate erythrocytes. I53-50 nanoparticles (NP) were prepared by incubation of domain 0-I53-50A (D0_FCoV-23), domain A-I53-50A, or domain B-I53-50A trimers with pentameric I53-50B at a 1:1 molar ratio, for at least 30 min at room temperature. Assembly of the CCoV-HuPn-2018 domain-0-I53-50 NP required mixing domain 0-I53-50A and bare I53-50A before adding pentameric I53-50B at 6 (1:5):1 molar ratio. **a,** Hemagglutination assay. **b,** SDS-PAGE analysis of purified nanoparticles. **c-f,** Representative electron micrographs of negatively stained FCoV-23 domain 0-(**c**), A-(**d**), or B-I53-50 NPs (**e**) and CCoV-HuPn-2018 domain 0-I53-50 NP (**f**). Scale bars, 200 nm. One hemagglutination assay corresponding to one preparation of nanoparticles is shown out of two biological replicates. **g,** Structure-based sequence alignment between FCoV-23 S and CCoV-HuPn-2018 S domain 0.

**Extended Data Figure 6.**
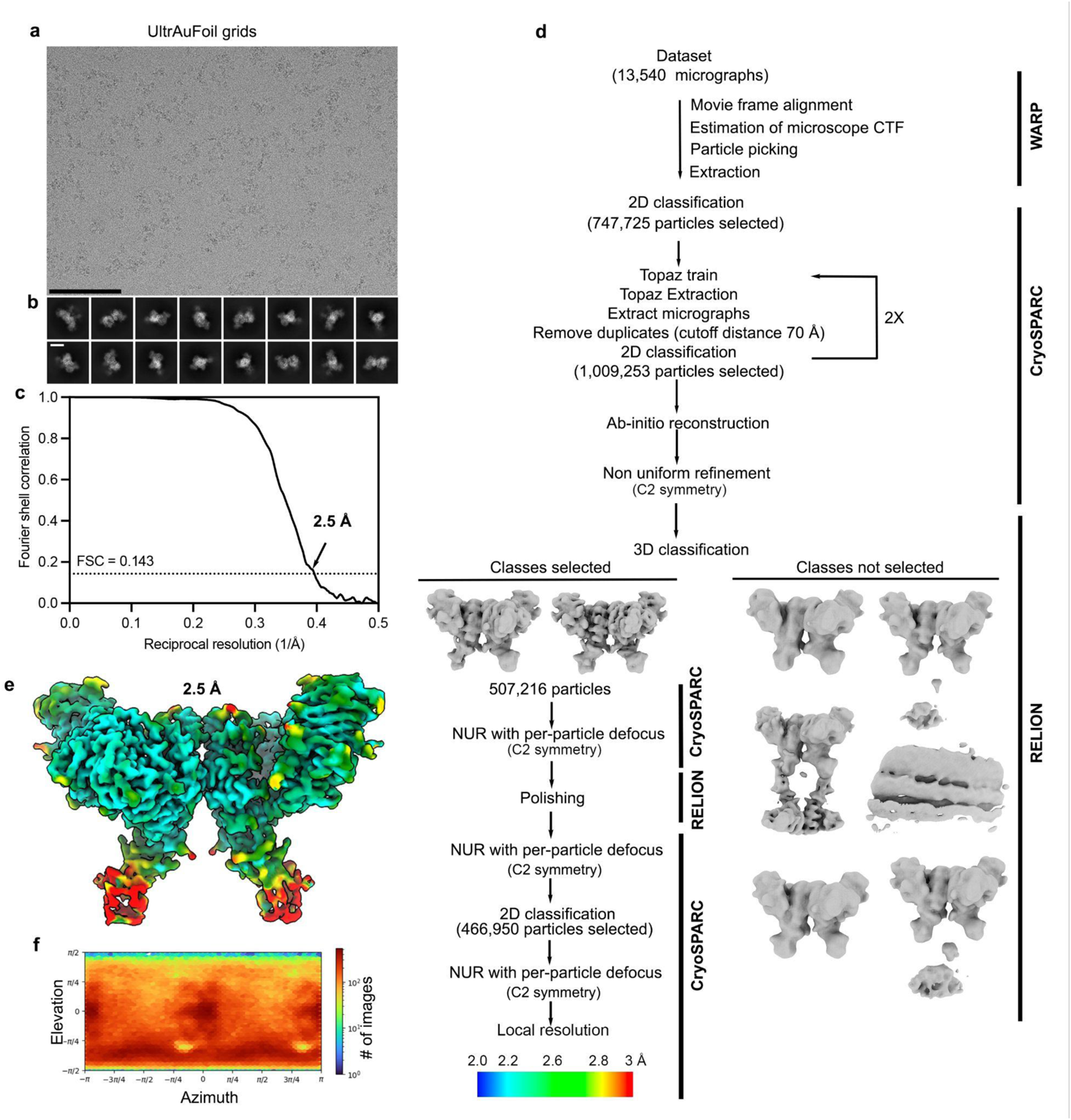
Data processing and validation of the FCoV-23 RBD in complex with feline APN. **a,b,** Representative electron micrograph (**a**) and 2D class averages (**b**) of the FCoV-23 RBD in complex with dimeric feline APN embedded in vitreous ice. Scale bar of the micrograph and the 2D class averages, 100 Å. (**c**) Gold-standard Fourier shell correlation curve for the FCoV-23 RBD-bound feline APN (solid black line). The 0.143 cutoff is indicated by a horizontal dotted gray line. (D) Cryo-EM data processing flowchart. (**e**) Unsharpened map of the FCoV-23 RBD-bound feline APN complex colored according to local resolution determined using cryoSPARC. (**f**) Angular distribution plots with all the particles contributing to the final map. CTF: contrast transfer function. NUR: non uniform refinement.

**Extended Data Figure 7.**
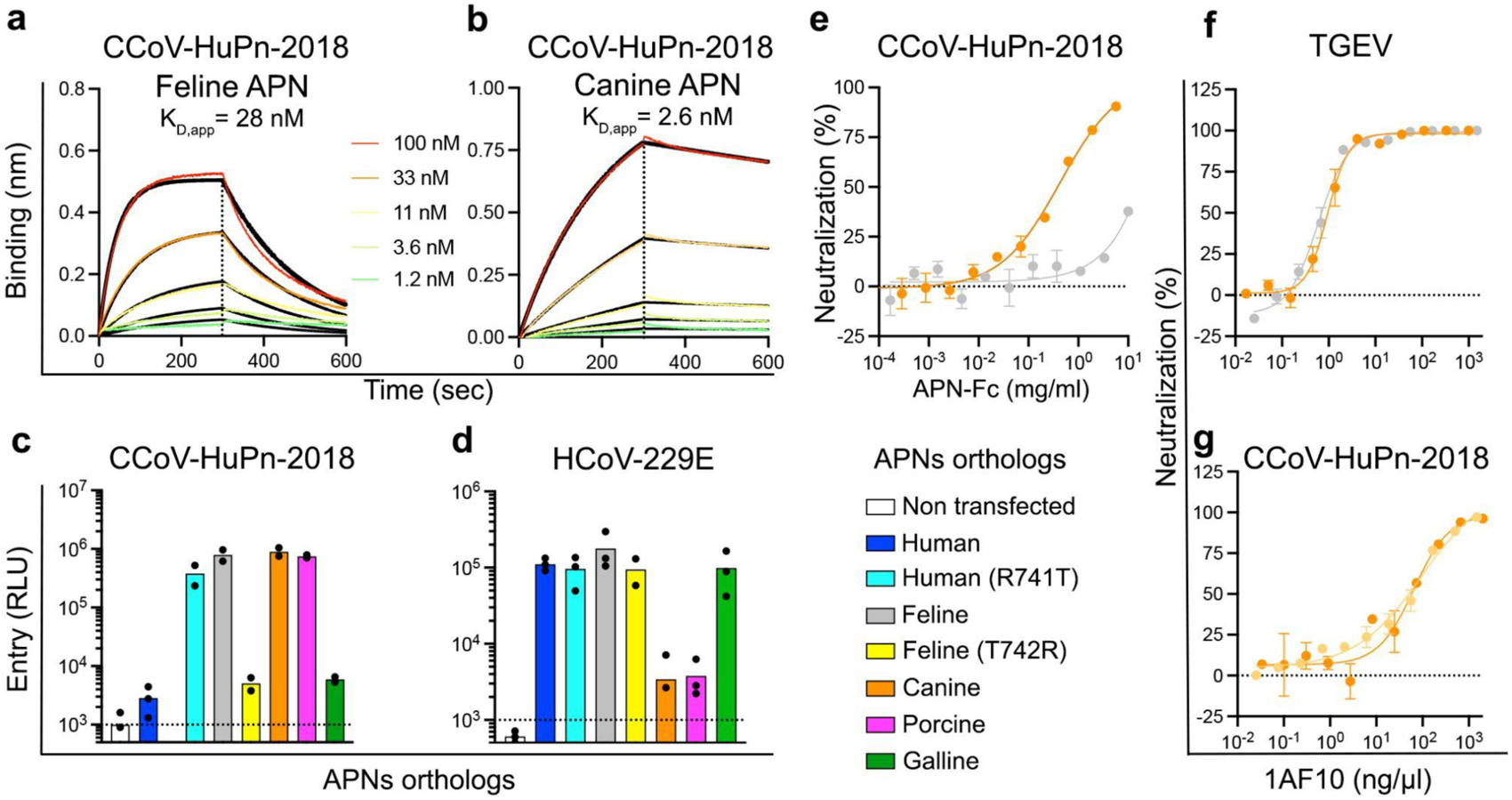
CCoV-HuPn-2018 and HCoV-229E usage of APNs orthologs. **a,b,** Biolayer interferometry binding kinetic analysis of the indicated concentrations of dimeric feline (**a**) and canine (**b**) APN-Fc ectodomains to the biotinylated CCoV-HuPn-2018 RBD immobilized on SA biosensors. Global fit (1:1 model) to the data is shown in black and reported affinities are expressed as apparent KD (KD, app) due to avidity resulting from the dimeric nature of APN. A representative experiment is shown out of two biological replicates. **c-d**, Entry of VSV particles pseudotyped with CCoV-HuPn-2018 S (**c**) or HCoV-229E S (**d**) into HEK293T cells transiently transfected with membrane-anchored human, human R741T (glycan knockin), feline, feline T742R (glycan knockout), canine, porcine and galline APN orthologs. RLUs, relative luciferase units. Each dot represents a biological experiment, each performed with technical duplicates or triplicates. **e**, Concentration-dependent inhibition mediated by purified dimeric feline and canine APN-Fc ectodomains of CCoV-HuPn-2018 S VSV pseudotyped virus entry into HEK293T cells transiently transfected with membrane-anchored canine APN. Curves represent one biological experiment performed with technical duplicates. Error bars represent the standard error of the mean (SEM). **f,g**, Dose-dependent neutralization of TGEV S (**f**) and CCoV-HuPn-2018 S (**g**) VSV pseudoviruses in the presence of various concentrations of the 1AF10 neutralizing monoclonal Fab fragment using HEK293T cells transiently transfected with membrane-anchored feline APN. Each curve represents one biological experiment, each performed with technical duplicates and the error bars represent the SEM.

**Extended Data Figure 8.**
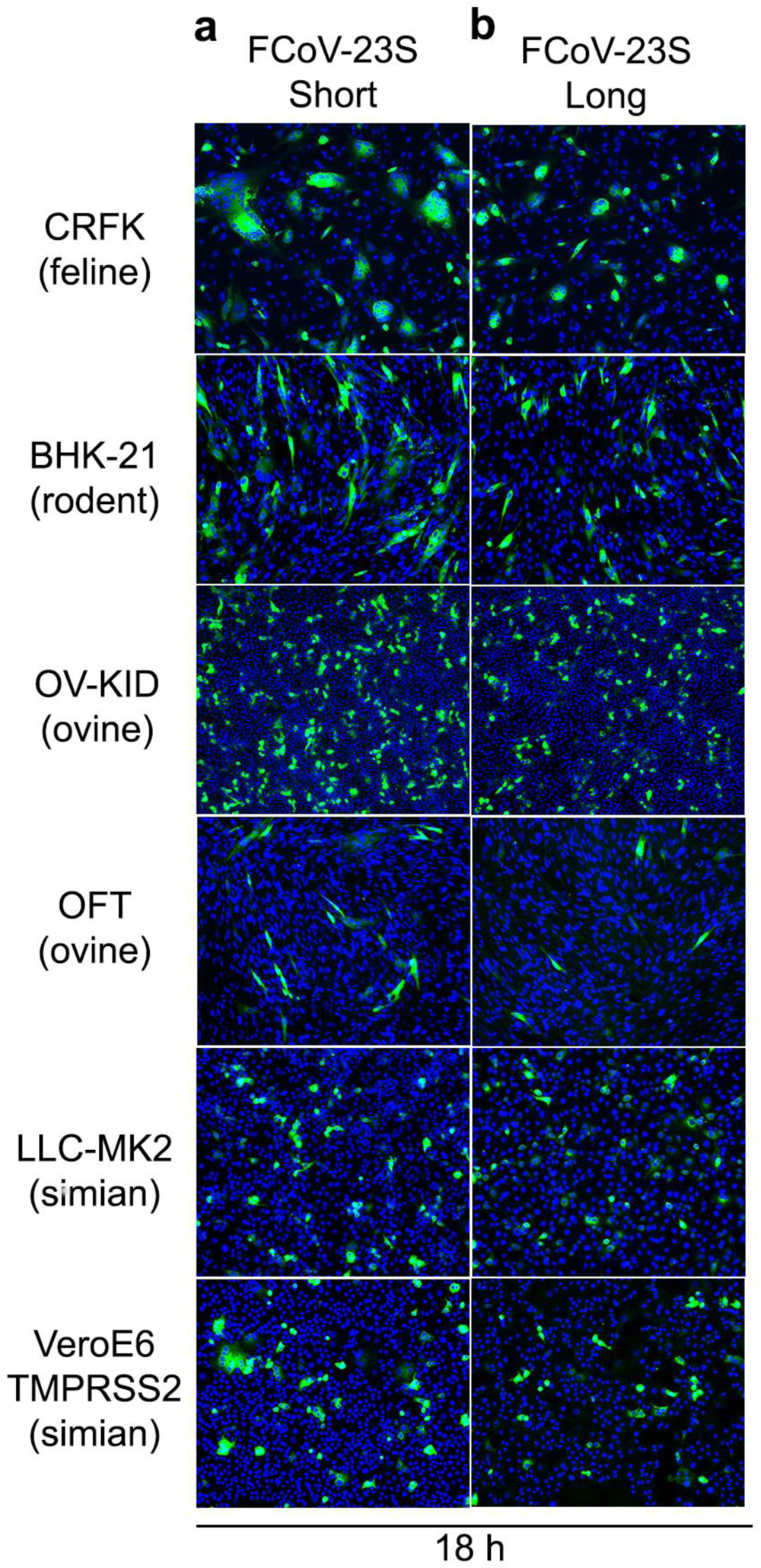
a, b, FCoV-23 S has limited or no permissiveness in rodent, ovine and simian cell lines. Cell-Cell fusion assay in the indicated cell lines expressing FCoV-23 S long or short S. Cells were fixed at 18 h post-transfection and FCoV-23 S long and short were detected using HA antibody (green) and nuclei were detected using DAPI stain (blue).

## Methods

### Cell lines

Cell lines used in this study obtained from ATCC were: human epithelial embryo cells (HEK293T, CRL-3216), human lung carcinoma epithelial cells (A549, CRM-CCL-185), human lung adenocarcinoma epithelial cells (Calu3, HTB-55), African green monkey kidney cells (VeroE6, CRL-1586), Rhesus monkey kidney cells (LLC-MK2, CCL-7), Syrian golden hamster kidney fibroblast cells (BHK-21, CCL-10), rat lung epithelial cells (L2, CCL-149), chicken embryo fibroblast cells (DF-1, CRL-3586), feline epithelial kidney cells (CRFK, CCL-94), feline airway epithelial cells (AK-D, CCL-150), canine tumor fibroblast cells (A-72, CRL-1542), canine epithelial kidney cells (MDCK, CCL-34), porcine epithelial kidney cells (LLC-PK1, CL-101) and bovine epithelial kidney cells (MDBK, CCL-22). Ovine kidney cells (OV KID) were from Cornell University Animal Health Diagnostic Center. Ovine fetal turbinate (OFT) cells were provided by Dr. Diego Diel. Feline macrophage-like cells, Fcwf-4, were initially obtained from ATCC (CRL-2787) and their progeny (Fcwf-CU) was selected by Dr. Edward Dubovi and Dr. Gary Whittaker at Cornell University to be significantly better at propagating feline coronavirus^82^. VeroE6 cells expressing transmembrane serine protease TMPRSS2 (VeroE6-TMPRSS2, JCRB1819) were obtained from the Japanese Collection of Research Bioresources (JCRB cell bank). Cell lines ExpiCHO cells and Expi293F cells were obtained from ThermoFisher Scientific. Cells were cultivated at 37°C, in the media recommended by the manufacturer, in an atmosphere of 5 % CO_2_ and with 130 rpm of agitation for suspension cells. Cell lines were not routinely tested for mycoplasma contamination.

### Plasmids

Genes used in this study were synthesized by GenScript, codon optimized for expression in mammalian cells, cloned into pcDNA3.1(+) between KpnI and XhoI, in frame with a Kozak’s sequence to direct translation, with the signal peptide derived from the µ-phosphatase: MGILPSPGMPALLSLVSLLSVLLMGCVAETGT (except for the full-length genes in which we used the original signal peptide), and avi tag and a C-terminal octa-histidine tag. FCoV-23 S long and short sequences were obtained from^22^. To stabilize FCoV-23 S ectodomain with domain 0 (S long) or without domain 0 (S short) in prefusion conformation, residues E (1146) and L (1147) were mutated to P, as previously described^44^.

Full-length wild-type S glycoproteins from HCoV-229E (AAK32191.1, P100E isolate) residues: 1-1,155, TGEV (ABG8935.1) residues: 1-1,425 and CCoV-HuPn-2018 (QVL91811.1) residues 1-1,425 harboring C-terminal deletions of 18-, 23- and 23-residues, respectively, were used to pseudotyped VSV(ΔG-luc). Full-length wild-type FCoV-23 S long and short also harbor a C-terminal deletion of 23 residues and the sequence GGS<u>YPYDVPDYA</u> was C-terminally incorporated (underlined is the HA-tag). FCoV-2023 S and CCoV-HuPn-2018 S B domains (also referred as receptor binding domains) matching with the full-length sequences indicated above, include residues: 529 to 677 and 523 to 671, respectively, and were fused to an avi-tag and his tag for biotinylation and affinity purification.

Full-length APNs were previously described^18^. Briefly, APNs from human (NP_001141.2), feline (NP_001009252.2), canine (NP_001139506.1), porcine (AGX93258.1) and galline (ACZ95799.1) orthologs comprise residues: 1-967, 1-967, 1-975, 1-963 and 1-967, respectively. APN ectodomains from human wildtype and R741T mutant (both comprising residues 66 to 967), feline (residues 64 to 967), canine (residues 71 to 975), porcine (residues 62 to 963) and galline (residues 69 to 967) from the same sequence codes shown above, were fused to a thrombin cleavage site followed by a human Fc fragment tag at the C-terminal end for affinity purification.

1AF10 Fab constructs for expression in mammalian cells of the light and heavy chain were previously described^18^. For nanoparticle assembly, FCoV-2023 domain 0 (residues 19-264), domain A (residues 265-504) and domain B (residues 529-677) were fused to the N-terminus of the trimeric I53-50A nanoparticle component^83^ using a 16 residue-long glycine/serine linker. CCoV-HuPn-2018 domain 0 (residues 17-259) fused to the N-terminus of the trimeric I53-50A nanoparticle component was previously described^18^.

### Mutagenesis

Full-length wild-type feline APN encoding plasmid was used as a template to knockout the glycan at position N740 (NWT to NWR) using the following not overlapping primers: 5’-aagaattgg<u>agg</u>gaccaccccc-3’ (underline is the codon change T to R) and 5-tgtcacgcgctcaaagtggttg-3’ using the fusion DNA polymerase (Thermofisher). After treating the PCR products with DpnI (New England Biolabs) during 1 h at 37°C, amplified plasmids were purified using PCR & DNA cleanup kit (Monarch) treated with T4 polynucleotide kinase (New England Biolabs) during 1 h at 37°C and ligated using T4 DNA ligase (New England Biolabs) at 25°C overnight before being used for transformation toOne Shot MAX Efficiency DH10B chemically competent cells (Invitrogen). Introduction of the desired mutations was verified by sequencing purified plasmids by Plasmidsaurus. Plasmid harboring the desired mutation was amplified and purified with EndoFree mega kit (Qiagen) to be suitable for transfection into mammalian cells.

### Protein expression and purification

To produce stabilized FCoV-23-S long and short and HCoV-229E S ectodomains, 200 ml of Expi293F cells grown to a density of 3 x 10^6^/mL and 37°C, were transfected with 640 µl of Expifectamine reagent (ThermoFisher) and 200 µg of the corresponding plasmids following manufacturer’s recommendations. The day after transfection, feed and enhancer were added to the cells and four to five days post-transfection, supernatants were clarified by centrifugation at 400 g for 15 minutes. Followed addition of 20 mM imidazole, 300 mM NaCl and 20 mM Tris-HCl pH 8.0, supernatants were further centrifuged at 14,000 g for 30 min and passed through a 1 mL Histrap Excel column (Cytiva) previously equilibrated with binding buffer (25 mM Tris pH 7.4 and 350 mM NaCl). FCoV-23 was eluted using a linear gradient of 500 mM imidazole. To express B domains from FCoV-23, CCoV-HuPn-2018 fused to an avi- and a histidine tag, 100 ml of Expi293F cells at 3 x 10^6^/mL were transiently transfected with 320 µl of Expifectamine and 100 µg of the respective plasmids, following the manufacturer’s indications. Four days post-transfection, supernatants were clarified by centrifugation at 800 g for 10 minutes, supplemented with 20 mM imidazole, 300 mM NaCl and 25 mM Tris-HCl pH 8.0, further centrifuged at 14,000 g for 30 min and passed through a 1 mL His trap HP column (Cytiva) previously equilibrated with binding buffer (25mM Tris pH 7.4 and 350mM NaCl). B domains were eluted using a linear gradient of 500 mM imidazole. A similar protocol was used to express and purify the Fab 1AF10 fused to 8 residues histidine tag except that 50 ml of Expi293F cells were transfected with a mixture containing: 50 µg of each individual plasmid encoding the Fab light and heavy chain and 160 µl of Expifectamine.

To express APN ectodomains from human, feline, canine, porcine and galline orthologs fused to Fc portion of human IgG, Expi293F cells were transiently transfected with the respective plasmids following the manufacturer’s protocols. Briefly, 50 ml of Expi293F cells at 3 x 10^6^/mL were transfected using 160 µl of Expifectamine and 50 µg of APN plasmid. Four days after transfection, supernatants were clarified by centrifugation at 800 g for 10 minutes, supplemented with 300 mM NaCl and 25 mM Tris-HCl pH 8.0, further centrifuged at 14,000 g for 30 min and passed through a 1 mL HiTrap Protein A HP column (Cytiva). Proteins were eluted using 0.1 M citric acid pH 3.0 in individual tubes containing 200 µl of 1 M Tris-HCl pH 9.0 to immediately neutralize the low pH needed for elution. APNs were further purified by size-exclusion chromatography (SEC) on a Superdex 200 column 10/300 GL (GE Life Sciences) previously equilibrated in 25 mM Tris pH 8.0 and 150 mM NaCl. Fractions containing the proteins were pooled and buffer exchanged to 25 mM Tris-HCl pH 8.0, 150 mM NaCl.

To produce feline APN without the Fc fusion, the Fc fragment was removed using thrombin (Millipore Sigma) in a reaction mixture containing: 3 µg of thrombin/mg of APN-Fc, 25 mM Tris-HCl pH 8.0, 150 mM NaCl and 2.5 mM CaCl_2_ incubated overnight at room temperature. The reaction mixture was loaded to a Protein A column to remove uncleaved APN-Fc and the Fc tag and feline APN was buffer-exchanged to 25 mM Tris-HCl pH 8.0, 150 mM NaCl.

To express 76E1 monoclonal antibody, Expi293F cells were transiently transfected with a plasmid encoding the full length IgG, following the manufacturer’s protocol. Four days after transfection, Expi293 cell supernatant was clarified by centrifugation at 4,121g for 30 minutes, supplemented with 20mM phosphate pH 8.0 and 0.5 mM PMSF. Supernatant was passed through a Protein A affinity column (Cytiva) twice before being washed with 20 mM phosphate pH 8.0. Antibody 76E1 was eluted with 100 mM citric acid pH 3.0 into tubes containing 1M Tris pH 9.0 for immediate neutralization of the low pH used for elution. Eluted 76E1 antibody was buffer-exchanged into 20 mM phosphate buffer pH 8.0/100 mM NaCl and purity was evaluated by SDS-PAGE.

### Protein Biotinylation

B domains of FCoV-CCoV S and CCoV-HuPn-2018 S were biotinylated using BirA biotin-protein ligase standard reaction kit (Avidity) following manufacturer’s protocol. In a typical reaction, 40 μM of B domains were incubated overnight at 4°C with 2.5 μg of BirA enzyme in reaction mixtures containing 1X BiomixB, 1X BiomixA and 40 µM BIO200. Domains B were further separated from Bir A by SEC using Superdex 75 increase 10/300 GL (GE LifeSciences) and concentrated using 10 kDa filters (Amicon).

### Biolayer interferometry (BLI) experiments

Biotinylated B domains from FCoV-23 S and CCoV-HuPn-2018 S were immobilized at 2 µg/mL in 10X kinetics buffer (Sartorius) to streptavidin (SA) biosensors (Sartorius) previously hydrated in 10X Kinetics Buffer for at least 10 minutes. Loaded tips were dipped into a solution containing 1 µM of APN orthologs from feline, canine, porcine, galline, human, and human R741T mutant in 10X Kinetics Buffer. For apparent K_D_ determination, loaded tips were dipped into various concentrations of feline or canine APN-Fc for 300 seconds followed by a 300 seconds dissociation phase in 10X Kinetics buffer. Data were baseline subtracted and the plots were fitted using the Sartorius analysis software (v.11.1). Data were plotted in Graphpad Prism (v.10.0.03). These experiments were done side-by-side with two different batches of B domains and APN orthologs preparations.

### Bacterial protein expression and purification of nanoparticles components

The I53-50A and I53-50B proteins were expressed as described before^83^. Briefly, transformed Lemo21(DE3) (NEB) in LB (10 g tryptone, 5 g yeast extract, 10 g NaCl) were grown at 37°C to an OD600 0.8 with agitation. Expression was induced with 1 mM IPTG and temperature was reduced to 18°C. Cells were harvested after 16 h and lysed by microfluidization using a Microfluidics M110P at 18,000 psi in 50 mM Tris, 500 mM NaCl, 30 mM imidazole, 1 mM PMSF, 0.75% CHAPS. Lysates were clarified by centrifugation at 24,000 g for 30 min and applied to a 2.6 3 10 cm Ni Sepharose 6 FF column (Cytiva) for purification by IMAC on an AKTA Avant150 FPLC system (Cytiva). Proteins were eluted with a linear gradient of 30 mM to 500 mM imidazole in 50 mM Tris pH 8,500 mM NaCl, 0.75% CHAPS buffer. Peak fractions were pooled, concentrated in 10K MWCO centrifugal filters (Millipore), sterile filtered (0.22 mm) and applied to either a Superdex 200 Increase 10/300, or HiLoad S200 pg GL SEC column (Cytiva) previously equilibrated in 50 mM Tris pH 8, 500 mM NaCl, 0.75% CHAPS buffer.

### *In vitro* nanoparticle assembly

Concentration of purified individual nanoparticle components was determined by measuring absorbance at 280 nm and the corresponding calculated extinction coefficients. Nanoparticles were prepared by incubation of domain 0-A-I53-50A, domain A-I53-50A or domain B-I53-50A trimers with pentameric I53-50B at molar ratios 1.1:1, respectively, in 50 mM Tris pH 8, 500 mM NaCl, 0.75% w/v CHAPS. Formation of CCoV-HuPn-2018 domain 0-NPs required mixing domain 0-I53-50A with I53-50A at a molar ratio of 1:6 with pentameric I53-50B. All *in vitro* assemblies were incubated at room temperature with gentle rocking for at least 20 min before subsequent purification by SEC on a Superose 6 column to remove residual unassembled components. Fractions were analyzed by negative stain electron microscopy and by PAGE-SDS. Assembled particles eluted in the void volume of a Superose 6 column and were pooled and stored at 4°C until use in hemagglutination assays.

### Negative stain electron microscopy

Nanoparticles diluted to 0.01 mg/mL in 50 mM Tris pH 8, 150 mM NaCl, were adsorbed to glow-discharged home-made carbon-coated copper grids for 30 seconds. The excess liquid was blotted away with filter paper (Whatman 1) and 3 µL of 2% w/v uranyl formate were applied to the grids. Finally, the stain was blotted away, and the grids were allowed to air dry for 1 min. Grids were imaged on a 120kV FEI Tecnai G2 Spirit with a Gatan Ultrascan 4000 4k x 4k CCD camera at 67,000 nominal magnification using a defocus ranging between 1.0 and 2.0 mm and a pixel size of 1.6 Å.

### Hemagglutination assay

The hemagglutination assay was performed according to standard procedures. Briefly, 25 µl of CCoV-HuPn-2018 domain 0, FCoV-23 domain 0, domain A, domain B NPs at 125 ng/ml were incubated with 25 µl of 1% feline (Cornell University), chicken (Lampire), human (Rockland Immunochemicals), bovine (Lampire) and turkey (Lampire) erythrocytes diluted in DPBS (Gibco) in V-bottom, 96-well plates (Greiner Bio-One) for 30 min at room temperature after which plates were photographed and hemagglutination was analyzed. CCoV-HuPn-2018 domain 0-NPs were used as a positive control.

### VSV pseudotyped virus production

FCoV-23, CCoV-HuPn, HCoV-229E (AAK32191.1, P100E isolate) and TGEV S pseudotyped VSV were generated as previously described^18^. Briefly, HEK293T cells in DMEM supplemented with 10% FBS and 1% PenStrep and seeded in poly-D-lysine coated 10-cm dishes were transfected with a mixture of 24 µg of the corresponding plasmid encoding for: CCoV-HuPn S, TGEV S or HCoV-229 S, 60 µl Lipofectamine 2000 (Life Technologies) in 3 ml of Opti-MEM, following manufacturer’s instructions. After 5 h at 37°C, DMEM supplemented with 20% FBS and 2% PenStrep was added. The next day, cells were washed three times with DMEM and were transduced with VSVΔG-luc^84^. After 2 h, virus inoculum was removed and cells were washed five times with DMEM prior to the addition of DMEM supplemented with anti-VSV-G antibody [Il-mouse hybridoma supernatant diluted 1 to 25 (v/v), from CRL-2700, ATCC] to minimize parental background. After 18-24 h, supernatants containing pseudotyped VSV were harvested, centrifuged at 2,000 x g for 5 minutes to remove cellular debris, filtered with a 0,45 µm membrane, concentrated 10 times using a 30 kDa cut off membrane (Amicon), aliquoted, and frozen at -80°C.

### MLV pseudotyped virus production

MLV pseudotyped particles were produced as previously described^85^. Briefly, HEK-293T cells were seeded at 5 x 10^5^ cells/ml in 6 well plates (Costar) a day before transfection. At ∼60% confluency, cells were transfected with 800 ng of pCMV-MLV gag-pol, 600 ng of pTG-Luciferase, and 600 ng of pCDNA3.1 FCoV-23 S short, pCDNA 3.1 FCoV-23 S long, pCAGGS empty vector as a negative control, or pCAGGS-VSV-G as a positive control and incubated at 34°C. Supernatants were collected 48 h post-transfection, cell debris was removed by centrifugation at 1000g and the supernatants-containing particles were filtered with a 0.45 µm membrane, aliquoted, and frozen at -80°C.

### Pseudotyped VSV infections and neutralizations

For pseudotyped VSV infections and neutralizations, HEK293T cells were transfected with plasmids encoding for the different full-length APN orthologs (flAPN) following a previously described protocol^18^. Briefly, HEK293T cells at 90% confluency and seeded in poly-D-lysine coated 10-cm dishes were transfected with a mixture in Opti-MEM containing 8-10 µg of the corresponding plasmid encoding flAPN orthologs and 30 µl of Lipofectamine 2000 (Life Technologies) according to the manufacturer’s instructions. After 5 h at 37°C, cells were trypsinized, seeded into poly-D-lysine coated clear bottom white walled 96-well plates at 50,000 cells/well and cultured overnight at 37°C. For infections, 20 µl of the corresponding pseudotyped VSV diluted 1 to 20 were mixed with 20 µl of DMEM and the mixture was added to the cells previously washed two times with DMEM. After 2 h at 37°C, 40 µl of DMEM was added and cells were further incubated overnight at 37°C.

For neutralizations, a single dilution of 1 to 10 for human or mice sera samples or eleven 3-fold serial dilutions of Fab 1AF10 or APN ectodomains, were prepared in DMEM. 20 µl of FCoV-23-, CCoV-HuPn-2018-, HCoV-229E- or TGEV-S pseudotyped VSV were added 1:1 (v/v) to each Fab1AF10, or APN ectodomains and mixtures were incubated for 45-60 min at 37°C. After removing media, transfected HEK293T cells were washed three times with DMEM and 40 μL of the mixture containing pseudotyped VSV:Fab/APN ectodomains/sera were added. One hour later, 40 μL DMEM were added to the cells. After 17-24 h, 60 μL of One-Glo-EX substrate (Promega) were added to each well and plates were placed in a shaker in the dark. After 5-15 min incubation, plates were read on a Biotek plate reader. Measurements were done in technical duplicates within at least two biological experiments. Relative luciferase units were plotted and normalized in Prism (GraphPad): cells alone without pseudotyped virus were defined as 0% infection, and cells with virus only (no sera) were defined as 100% infection. Most of the human sera was collected from prospective bone marrow donors in Seattle with approval from the Human Subjects Institutional Review Board in the 1980s and were stored in the Infectious Disease Sciences Biospecimen Repository at the Vaccine and Infectious Disease Division of the Fred Hutch Cancer Center.

### Pseudotyped MLV infection

Fcwf-CU, CRFK, AK-D, A-72, MDCK, LLC-MK2, VeroE6, VeroE6-TMPRSS2, A549, Calu3, LLC-PK1, MDBK, OFT, OVKID, L2, BHK 21, and DF-1 cells were seeded at 3-5 x 10^5^ cells/mL in a 24-well plate (Costar) and cultured at 37°C. At ∼90-100% confluency, cells were infected with 200 µL of MLV particles and incubated on a rocker at 37°C. After 1.5 hours, 300 µL of the corresponding complete media was added to cells. After 72 h, cells were lysed with 100 µL of 1X luciferase cell culture lysis reagent (25 mM Tris-phosphate pH 7.8, 2 mM DTT, 2 mM 1,2- diaminocyclohexane-N,N,Ń,Ń-tetraacetic acid, 10% glycerol, 1% Triton X-100, Promega), and luminescence was measured using the Luciferase Assay System (Promega). Luciferase activity was measured by adding 20 µL of luciferin substrate to 10 µL of cell lysate, using a GloMax 20/20 luminometer (Promega). MLV infection assays were performed in triplicate.

### Fluorogenic peptide cleavage assay

FCoV-23 S_2’_ fluorogenic peptide, _958_SKRKYRSAIE_967_, site was synthesized by Biomatik with a 7-methoxycoumarin-4-yl acetyl (MCA) group covalently bound at the N-terminus end and a 2,4-dinitrophenyl (DNP) group at the C-terminal end was used as a substrate in a fluorogenic peptide cleavage assay performed as described previously^86^. Briefly, FCoV-23 S_2’_ peptide was resuspended in water to 1mM and cleavage was performed in a black, flat-bottom 96-well plate (Costar) in a total volume of 100 µL/well as follow: 94.5 µL of enzyme-specific buffer, 5 µL of S_2’_ fluorogenic peptide was added to each well (50 µM/well), and 0.5 µL of protease. TPCK trypsin (Thermofisher Scientific) was used at 4.3 nM/well in 67 mM NaH_2_PO_4_ pH 7.6. Furin (NEB) was used at 1 U/well in 20 mM HEPES-NaOH pH 7.5, 0.2 mM CaCl_2_, and 1 mM 2-mercaptoethanol. PC1 (NEB) was used at 1 U/well in 100 mM HEPES-NaOH pH 6.0, 1 mM CaCl_2_, and 1 mM 2-mercaptoethanol. Xa (NEB) was used at 1 U/well in 20 mM Tris-HCl pH 8.0, 100 mM NaCl, and 2 mM CaCl_2_. Plasmin (Innovative Research) was used at 0.5 µg/mL/well in 100 mM Tris-HCl pH 7.4, 0.1% Tween 20, and 0.1 mM EDTA. Cathepsin L (R&D) was used at 0.5 µg/mL/well in 50 mM MES-NaOH pH 6 , 5 mM DTT, 1 mM EDTA, and 0.005% Brij-35. After buffer, protease, and peptide were mixed, fluorescence was measured every 60 seconds for 2 h at 37°C using the SpectraMax Gemini XPS (Molecular Devices) at an excitation wavelength of 330 nm and an emission wavelength of 390 nm. Data were plotted with relative fluorescent units (RFU) on the y-axis and time on the x-axis. Vmax was calculated by determining the slope of the linear range.

### Western blot

For VSV pseudotypes characterization, 15 µl of each pseudotype were mixed with 4X SDS-PAGE loading buffer, run through a 4%–15% gradient Tris-Glycine gel (BioRad) and transferred to a PVDF membrane using the protocol mix molecular weight of the Trans-Blot Turbo System (BioRad). The membrane was blocked with 5% milk in TBS-T (20 mM Tris-HCl pH 8.0, 150 mM NaCl) supplemented with 0.05% Tween-20 at room temperature and with agitation. After 1 h, the fusion-peptide-specific 76E1 monoclonal antibody was added at 2 µg/ml and incubated overnight at 4°C with agitation. Next day, the membrane was washed three times with TBS-T and an Alexa Fluor 680-conjugated goat anti-human secondary antibody (1:50,000 dilution, Jackson ImmunoResearch, 109-625-098) was added and incubated during 1 h at room temperature. Membrane was washed three times with TBS-T after which a LI-COR processor was used to develop the western blot.

### Cell-Cell fusion assay

FCWF-4-CU, CRFK, AK-D, A-72, Vero E6, A549, BHK, and OV KID cells were seeded at 3-5 x 10^5^ cells/mL in a 24-well plate (Costar) and cultured at 37°C. Cells were transfected at 80% confluency with 250 ng of pCDNA3.1 vector encoding FCoV-23 S short or FCoV-23 S long using Lipofectamine 3000 (Thermofisher Scientific). After 4 to 18 hours post-transfection, cells were fixed with 4% paraformaldehyde, for 20 minutes at room temperature, washed 3 times with PBS and permeabilized with 0.1% Triton X-100 for 10 minutes. After 3 more washes with PBS, cells were incubated with anti-HA antibody (Thermofisher Scientific) at 1/300 dilution in 1% Bovine Albumin (VWR) in PBS for 3 h at room temperature. Secondary antibody labeling was performed using AlexaFluor 488 goat-anti-mouse IgG (Invitrogen) at 1/300 dilution in 1% Bovine Albumin (VWR) in PBS during 45 min. After 3 final washes with PBS, nuclei were stained with DAPI Fluoromount-G (Southern Biotech) diluted 1/10 in PBS, and the cells were imaged at 10X magnification using an ECHO Revolve fluorescence microscope.

### CryoEM sample preparation, data collection and data processing for FCoV-23 S short and long

3 µl of FCoV-CCoV S without domain 0 (short) at approximately 0.1-0.3 mg/ml were loaded three times onto freshly glow-discharged NiTi grids covered with a thin layer of home-made continuous carbon prior to plunge freezing using a vitrobot MarkIV (ThermoFisher Scientific) with a blot force of -1 and 4.5 sec blot time at 100% humidity and 21°C. For FCoV-CCoV S with domain 0 (long), 3 µl of S at 0.15 mg/ml were loaded onto lacey grids freshly glow-discharged and covered with a thin layer of home-made continuous carbon following the same protocol as for FCoV-CCoV S short.

For FCoV-CCoV S short and long, 11,208 and 13,099 movies were collected, respectively, with a defocus range between -0.8 and -2.0 μm. Both datasets were acquired using a FEI Titan Krios transmission electron microscope operated at 300 kV equipped with a Gatan K3 direct detector and a Gatan Quantum GIF energy filter, operated with a slit width of 20eV. Automated data collection was carried out using Leginon software^87^ at a nominal magnification of 105,000x, pixel size of 0.843 Å. The dose rate was adjusted to 9 counts/pixel/s, and each movie was acquired in counting mode fractionated in 100 frames of 40 ms. Movie frame alignment, estimation of the microscope contrast-transfer function parameters, particle picking and extraction were carried out using Warp^88^. One round of reference-free 2D classification was performed using cryoSPARC^89^ with binned particles to select well-defined particle images. To further improve particle picking we trained Topaz picker^90^ on the Warp-picked particles on the selected classes after 2D classification. Topaz-picked particles were extracted and 2D classified using cryoSPARC. Topaz-duplicated picked particles were removed using 60 Å (for FCoV-23 S short) or 90 Å (for FCoV-23 S long) as a minimum distance cutoff. Initial model generation was done using ab-initio reconstruction in cryoSPARC and used as reference for a non-uniform refinement^91^ (NUR) (for FCoV-23 S short) or heterogenous refinement followed by NUR (for FCoV-23 S with domain 0) in cryoSPARC allowing the identification of two conformations for the FCoV-23 S long dataset. Particles were transferred from cryoSPARC to Relion^92^ using pyem^93^ (https://github.com/asarnow/pyem) to be subjected to one round of 3D classification with 50 iterations, using the NUR map as a reference model (angular sampling 7.5 ° for 25 iterations and 1.8 ° with local search for 25 iterations) and without imposing symmetry. Selected particles were subjected to a NUR using cryoSPARC. Particles were subjected to Bayesian polishing^94^ using Relion during which were re-extracted with a box size of 512 pixels at a pixel size of 1 Å. Another round of 2D classification was performed in cryoSPARC which was followed by a NUR including per-particle defocus refinement.

To improve the density of the domain 0 from the FCoV-23 S long and short map, particles were symmetry-expanded (for the FCoV-23 S long map) or not (FCoV-23 S short map) and subjected to a Relion 3D focus classification without refining angles and shifts using soft masks encompassing domain 0 in “swung-out” conformation from the FCoV-23 S long map or domain 0 in “proximal” conformation from the FCoV-23 S short map. Local refinement and local resolution estimation were carried out using CryoSPARC. Reported resolutions are based on the gold-standard Fourier shell correlation using 0.143 criterion^95^ and Fourier shell correlation curves were corrected for the effects of soft masking by high-resolution noise substitution^96^.

### CryoEM sample preparation, data collection and data processing for FCoV-RBD in complex with feline APN

Purified feline APN was incubated overnight at 4°C with a molar excess of purified feline RBD of 1:3 and the complex was purified by SEC. 2.7 µl of feline APN/feline RBD at 4 or 6 mg/ml was applied for 15 sec onto freshly glow discharged UltrAuFoil R 2/2 grids with 0.3 µl of 30 or 60 mM of CHAPSO, respectively, prior to plunge freezing using a vitrobot MarkIV (ThermoFisher Scientific) with a blot force of -0 and 6 sec blot time at 100% humidity and 21°C. 13,540 micrographs were collected using Leginon^87^, with a defocus range between -0.8 and - 2.0 μm on the same microscope set up described for the FCoV-23 S glycoproteins ectodomain. Data processing was similar as described for FCoV-23 S glycoproteins ectodomain except that 2 cycles of Topaz picker^90^ on the Warp-picked particles on the selected classes after 2D classification was performed.

### CryoEM model building and analysis

Model Angelo^97^ was used to generate an initial model and UCSF Chimera^98^ and Coot^99^ were used to manually build the model. Model was refined and rebuilt into the maps using Coot and Rosetta^100,101^. Model validation was done using Molprobity^102^ and Privateer^103^. Figures were generated using UCSF ChimeraX^104^.

### In vivo immunogenicity Study

Female BALB/cAnNHsd mice were purchased from Envigo (order code 047) at 7 weeks of age and were maintained in a specific pathogen-free facility within the Department of Comparative Medicine at the University of Washington, Seattle, accredited by the Association for Assessment and Accreditation of Laboratory Animal Care (AAALAC). Animal experiments were conducted in accordance with the University of Washington’s Institutional Animal Care and Use Committee under protocol 4470-01. For each immunization, low endotoxin HCoV-229E S ectodomain was diluted to 100 µg/mL in buffer and mixed 1:1 (v/v) with AddaVax adjuvant (InvivoGen vac-adx-10) to obtain a final dose of 5 µg of HCoV-229E S ectodomain per animal, per injection. At 8 weeks of age, 10 mice per group were injected subcutaneously in the inguinal region with 100 μL of immunogen at weeks 0, 3, and 6. Animals were bled via submental venous puncture at weeks 2 and 5 and terminal blood was collected at week 8. Whole blood was collected in serum serum separator tubes (BD #365967) and rested at room temperature for 30 min for coagulation. Tubes were then centrifuged for 10 min at 2,000 x *g* and serum was collected and stored at -80°C until use.

